# CRF release from a unique subpopulation of accumbal neurons constrains action-outcome acquisition in reward learning

**DOI:** 10.1101/2023.11.16.567495

**Authors:** Elizabeth A. Eckenwiler, Anna E. Ingebretson, Jeffrey J. Stolley, Maxine A. Fusaro, Alyssa M. Romportl, Jack M. Ross, Christopher L. Petersen, Eera M. Kale, Michael S. Clark, Selena S. Schattauer, Larry S. Zweifel, Julia C. Lemos

**Author notes:** Corresponding author: Dr. Julia C. Lemos, 321 Church Street, Minneapolis, MN 55406.

## Abstract

**Background:** The nucleus accumbens (NAc) mediates reward learning and motivation. Despite an abundance of neuropeptides, peptidergic neurotransmission from the NAc has not been integrated into current models of reward learning. The existence of a sparse population of neurons containing corticotropin releasing factor (CRF) has been previously documented. Here we provide a comprehensive analysis of their identity and functional role in shaping reward learning.

**Methods:** To do this, we took a multidisciplinary approach that included florescent in situ hybridization (N_mice_ ≥ 3), tract tracing (N_mice_ = 5), ex vivo electrophysiology (N_cells_ ≥ 30), in vivo calcium imaging with fiber photometry (N_mice_ ≥ 4) and use of viral strategies in transgenic lines to selectively delete CRF peptide from NAc neurons (N_mice_ ≥ 4). Behaviors used were instrumental learning, sucrose preference and spontaneous exploration in an open field.

**Results:** Here we show that the vast majority of NAc CRF-containing (NAc^CRF^) neurons are spiny projection neurons (SPNs) comprised of dopamine D1-, D2- or D1/D2-containing SPNs that primarily project and connect to the ventral pallidum and to a lesser extent the ventral midbrain. As a population, they display mature and immature SPN firing properties. We demonstrate that NAc^CRF^ neurons track reward outcomes during operant reward learning and that CRF release from these neurons acts to constrain initial acquisition of action-outcome learning, and at the same time facilitates flexibility in the face of changing contingencies.

**Conclusion:** We conclude that CRF release from this sparse population of SPNs is critical for reward learning under normal conditions.

## Introduction

The nucleus accumbens (NAc) drives several evolutionarily conserved behaviors, including environmental exploration and reward learning (1, 2). It is a part of the ventral striatum that interfaces with several different glutamatergic, monoaminergic and peptidergic inputs to influence motivated behaviors (3, 4). Based on groundwork done in the dorsal striatum, the NAc has been shown to similarly contain two populations of spiny projection neurons (SPNs) that can be distinguished by their peptidergic expression, intrinsic excitability, dopamine receptor expression and projection patterns (3, 5). Synchronous optogenetic stimulation of these distinct SPN populations often results in opposite behavioral responses — though this canonical behavioral pattern has recently been challenged (6). The accumbens also contains several interneuron subtypes, including cholinergic interneurons and several distinct GABAergic interneurons (3). Advancements in single cell (single nucleus) RNA-sequencing technologies has revealed a further diversity of cell populations that can be clustered in novel ways compared to traditional classification (7–9). It has also revealed an abundance of neuropeptides whose neurotransmission has not been integrated into behavioral models of NAc function (7).

A sparse population of neurons containing the neuropeptide neurotransmitter corticotropin-releasing factor (CRF) (10) was initially identified in the rat NAc and later confirmed in mice (11). These accumbal CRF neurons have been presumed to be a type of interneuron, primarily due to their scarcity. Yet, optogenetic stimulation of this population increases cFos activity in other brain regions such as the ventral tegmental area, ventral pallidum and bed nucleus of the stria terminalis (BNST), in addition to the NAc, leaving open the possibility for long-range projections (12). CRF has historically been characterized as a peptide that encodes the aversive properties of stress. However, within the corticomesolimbic pathway it may play a more complex role, such as the invigoration of exploratory and appetitive behaviors (13–15). In 2021, Baumgartner and colleagues demonstrated that optogenetic stimulation of NAc^CRF^ neurons in the rat promoted reward consumption and supported intracranial self-stimulation by increasing incentive motivation, not by producing aversion (12). To our knowledge, this was the first behavior linked to NAc^CRF^ neurons, yet it was unclear if these results were due to CRF release itself.

In this study, we thoroughly characterize NAc^CRF^ neurons and demonstrate a behavioral role for this population in instrumental learning. A combination of in situ hybridization, viral tract tracing and electrophysiology led us to conclude that NAc^CRF^ neurons are a unique subpopulation of SPNs that primarily project to the ventral pallidum and are, as a population, more excitable than SPNs. We find that NAc^CRF^ neurons increase their activity during reward seeking and that this activity scales depending on reward delivery. Importantly, CRF deletion from accumbal neurons facilitates faster acquisition of action-outcome reward learning and impairs reversal learning. Together, these results provide the first in-depth transcriptional, anatomical, and electrophysiological analysis of NAc^CRF^ neurons and identifies a critical role for accumbal CRF release in reward learning.

## Methods

For detailed methods, see *Supplemental Methods*.

### Animals

All procedures were done in accordance with the University of Minnesota Animal Care and Use Committee. Male and female were used in approximately equal numbers. We used mice from the C57BL/6J (JAX stock: 000664) and *Crh*^IRES-Cre/-^ (012704) lines. In some cases, *Crh^IRES-Cre/-^* mice were crossed with Ai14 mice (007914). Our validation of *Crh^IRES-Cre/-^* mice demonstrated that the specificity of the Cre-recombinase to NAc^CRF^ neurons was very high (>85%). However, the penetrance of Cre was significantly lower in NAc (38 ± 4%) compared to the paraventricular nucleus, BNST and central amygdala (Figure S1). For conditional knockout experiments, *Crh*^loxP/loxP^ mice generated by Dr. Michael S. Clark and validated by Dr. Larry Zweifel at the University of Washington were used (16).

### Fluorescent in situ hybridization and image analysis

RNAscope manual fluorescent multiplex (v1,v2) and HiPlex assays were run as detailed in ACD technical manuals and previously reported (17). Tissue was imaged on Nikon AR-Flim or Leica Stellaris 8 confocal microscopes (20x tiled). For each run of RNAscope, capture settings were identical across animals. For analysis, HALO image analysis platform (Indica Labs) or Fiji/ImageJ was used.

### Immunohistochemistry and image analysis

Mice were perfused, tissue prepared, and immunohistochemistry performed as previously described (18). For tract tracing studies, images were acquired using a confocal microscope (Stellaris 8, Leica). Three to five confocal images were acquired per region per mouse, with 5 mice total. Analysis was performed using ImageJ software.

### Surgical procedures

For detailed surgical procedures, see *Supplemental Methods*. Adult mice were quickly and deeply anesthetized using isoflurane (4% induction, 2% maintenance), and the skull leveled on a stereotaxic frame (Kopf, Model 1900). Viral particles were delivered using handmade 30g metal injectors connected to a 2µL syringe (Hamilton) with polyethylene tubing and delivered via pump (WPI microinjection syringe pump) at 100nL/min. For some experiments, we also use glass sharp electrodes and Drummond Nanoject II microinfusion system. For NAc targeting, the following coordinates were used: AP = 1.15/1.2, ML = ±1.1, DV = -4.7/4.75; mm relative to bregma. For fiber photometry experiments, NAc coordinates were: AP = 1.3, ML = ±1.3, DV = -4.7. For VP targeting: AP= 0.3, ML: ±1.1, DV -5.0.

### Ex vivo electrophysiology

Ex vivo electrophysiology techniques were conducted as previously described (18). For optogenetic experiments, sections from *Crh*^IRES-Cre/-^ mice injected into NAc with a Cre-dependent channelrhodopsin-expressing virus were prepared for either midbrain or ventral pallidum recordings. Paired pulse optogenetic stimulation was produced by a 470nm LED (M470F3, ThorLabs) connected to a T-cube driver (LEDD1B, Thorlabs).

### Operant training

Fixed ratio training was done as previously described(19), with slight modifications (see *supplemental material*). Progressive ratio tasks and reversal tasks were done as previously described (20),(21).

### Fiber photometry

During behavioral training, mice were attached to a fiberoptic patchcord (Doric) every day, but recordings were only performed on alternating days to prevent bleaching. Analysis of photometry signals was done using the Python toolbox GuPPy, created by the Lerner Lab (22). An isosbestic control channel (405 nm) was used to filter out artifacts, and a standard z-score was calculated. Med-PC-generated timestamps were used to align z-scores from behavioral events. Z-scores were first averaged by session then by mouse.

## Results

### Determination of NAc^CRF^ neuron transcriptomic features

Using fluorescent in situ hybridization in tissue from male and female C57BL6/J, we determined co-expression distribution of *Crh* mRNA with markers for SPNs and interneurons. Collectively, these markers cover the preponderance of classically defined neuronal types in the NAc. *Crh* mRNAs were present in 2.5% of NAc cells and 0.54% of dorsal striatal cells, with a significant low-to-high rostral-caudal gradient in *Crh*+ expression (Figure S2). The majority of *Crh*+ neurons expressed *Drd1* (66 ± 6%) or *Pdyn* (74 ± 3%), canonical markers for direct pathway SPNs. A smaller proportion of *Crh+* neurons expressed *Drd2* (36 ± 5%) or *Penk* (24 ± 3%), canonical indirect pathway markers (Figure 1a,b). *Crh*+ neurons constituted 3.0 ± 0.8% of all *Drd1+* neurons and 1.2 ± 1.0% of all *Drd2*+ neurons (Figure 1c). We found minimal co-expression of *Crh* with most striatal interneuron markers (Figure 1d). However, there were small but appreciable percentages of co-expression of *Crh* with *Calb2* (9.0 ± 1.2%) and *Pthlh* (7.5% ± 1.2%) (Figure 1d). When assessing traditional SPN markers using HiPlex rather than MultiPlex methods, we found the majority of *Crh+* neurons expressed *Drd1+* exclusively (51.1±1.6%) (Figure 1f). Approximately half of *Crh+/Drd2+* cells did so exclusively (10.6 ± 0.6%), while the other half also expressed *Drd1* (11.0 ± 1.7%) (Figure 1f).

**Figure 1.**
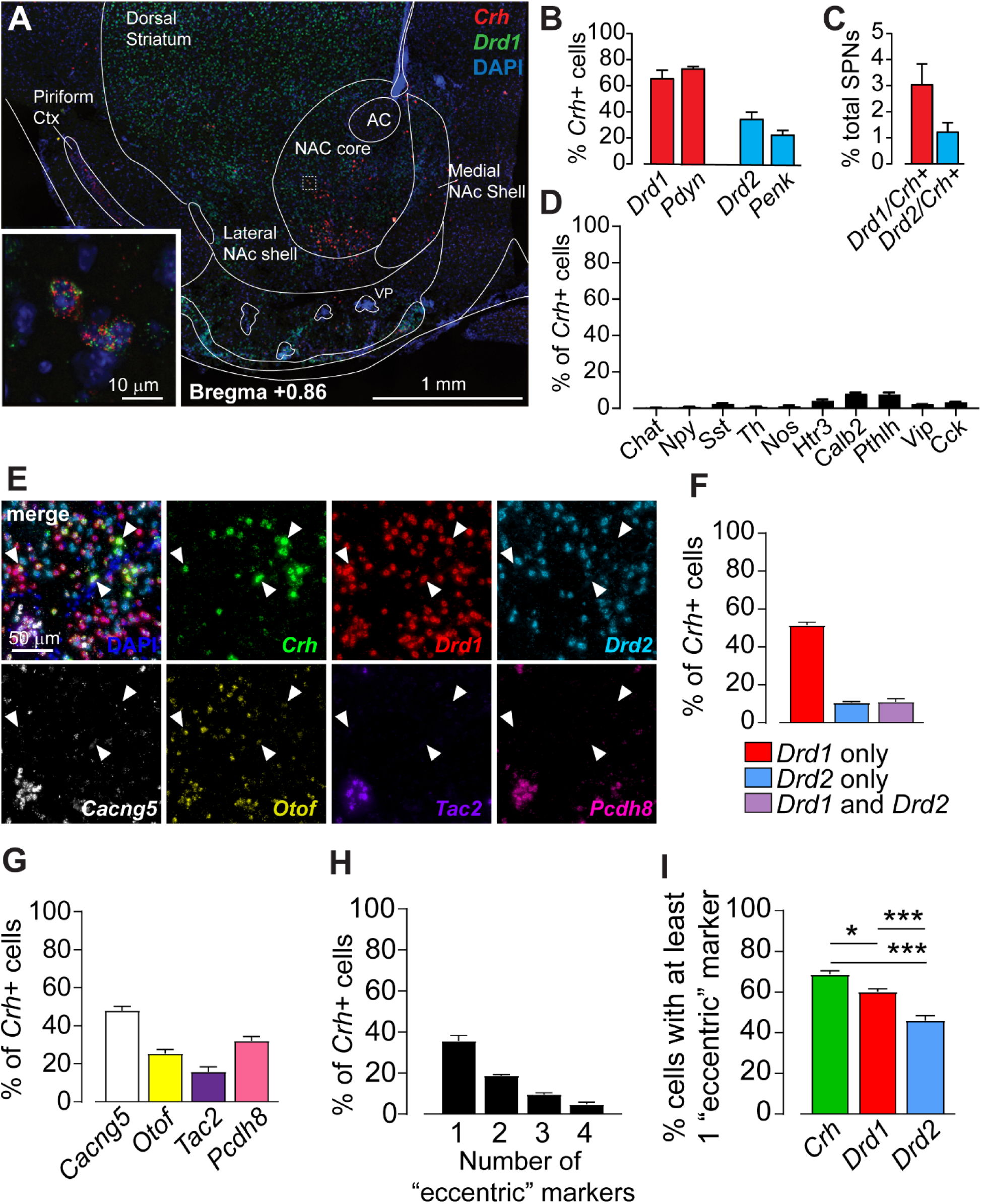
Characterization of NAc CRF neurons using fluorescent in situ hybridization: (A) Representative image of fluorescent in situ hybridization expression in the NAc and co-expression of *Crh* (red) and *Drd1* (green) mRNA counterstained with DAPI (blue). (B) Quantification of co-localization of *Drd1* (N = 12), *Pdyn* (N = 3), *Drd2* (N = 11), and *Penk* (N = 3) mRNA transcripts in *Crh*-expressing cells. (C) Proportion of *Drd1* (N = 12) and *Drd2* (N = 11) populations expressing *Crh* mRNA. (D) mRNA expression of interneuron markers in *Crh*+ cells: *Chat* (0.2 ± 0.1%, N = 4)*, Npy* (0.6 ± 0.2%, N = 10)*, Sst* (2.3 ± 0.5%, N = 10)*, Th* (0.7 ± 0.3%, N = 4)*, Nos* (1.0 ± 0.6%, N = 4)*, Htr3* (2.5 ± 0.6%, N = 6)*, Calb2* (9.0 ± 1.2%, N = 8)*, Pthlh* (7.5% ± 1.2%, N = 7)*, Vip* (1.2 ± 0.1%, N = 5), and *Cck* (3.2 ± 0.4%, N = 7) (E) Representative images from the HiPlex assay, demonstrating fluorescent labeling of seven distinct mRNA transcripts in the same tissue, counterstained with DAPI. To characterize the *Crh*+ population, probes for *Crh*, canonical SPN markers (*Drd1*: red; *Drd2*: blue), and “eccentric” SPN markers (*Cacng5*: white; *Otof*: yellow; *Tac2*: purple; *Pcdh8*: pink) were simultaneously visualized in the NAc. (F) Quantification of the HiPlex assay for *Drd1* only*, Drd2* only or *Drd1/Drd2* co-expressing cells within the *Crh* population (N = 8). (G) Analysis of co-localization of “eccentric” markers (*Cacng5, Otof, Tac2,* and *Pcdh8*) with *Crh* mRNA (N = 8). (H) Distribution of *Crh* cells expressing either one, two, three or all four eccentric markers assayed (I) Comparison of the proportion of *Crh, Drd1,* or *Drd2* cells expressing at least one “eccentric” marker. The *Crh* population had a significantly higher proportion of cells expressing at least one “eccentric” marker compared to *Drd1* and *Drd2* populations. Additionally, a higher proportion of *Drd1* cells expressed “eccentric” markers compared to *Drd2* cells (*p = 0.0114, ***p < 0.0001, one-way ANOVA with Sidak post-hoc unless otherwise noted; N = 8). (Data are represented as mean ± SEM. All N represent the number of animals used.)

Saunders et al., (2018) identified a third cluster of “eccentric” SPNs in the dorsal striatum that could be distinguished by the presence of one of four markers (*Otof, Tac2, Pcdh8* and *Cacng5*)(23). Using HiPlex techniques, we found that 69 ± 2% of *Crh+* cells contained at least one of these markers (Figure 1g,h), compared to 60 ± 1% of all *Drd1+* neurons and 46 ± 2% of all *Drd2+* neurons (one-way ANOVA with Sidak; main effect F_2,21_ = 37.91, p < 0.0001). These data indicate that, despite being a relatively small population, CRF cells overrepresent “eccentric” markers compared to SPN populations overall.

### NAc^CRF^ neurons project to ventral pallidal and midbrain regions

We next transduced the NAc of *Crh*^IRES-Cre/-^ mice bilaterally with a Cre-dependent virus expressing GFP in transfected cells and along fibers of passage, with mRuby tagged to synaptophysin and labeling presynaptic boutons. We used tyrosine hydroxylase immunoreactivity to identify midbrain dopamine neurons and substance P to delineate the ventral pallidum (VP) for quantification. Synaptic boutons in the VP and in the substantia nigra *pars compacta/reticulata* were detected based on the presence of GFP+ fibers and mRuby+ puncta (Figure 2a-c). We found significantly more mRuby+ puncta in the VP compared to the midbrain (t-test, p = 0.0165) (Figure 2d). Interestingly, we found evidence for mRuby+ varicosities along CRF-containing SPNs, indicating *en passant* axonal release sites, and several instances of NAc^CRF^ clusters at the border between the NAc and the VP (Figure S3).

**Figure 2.**
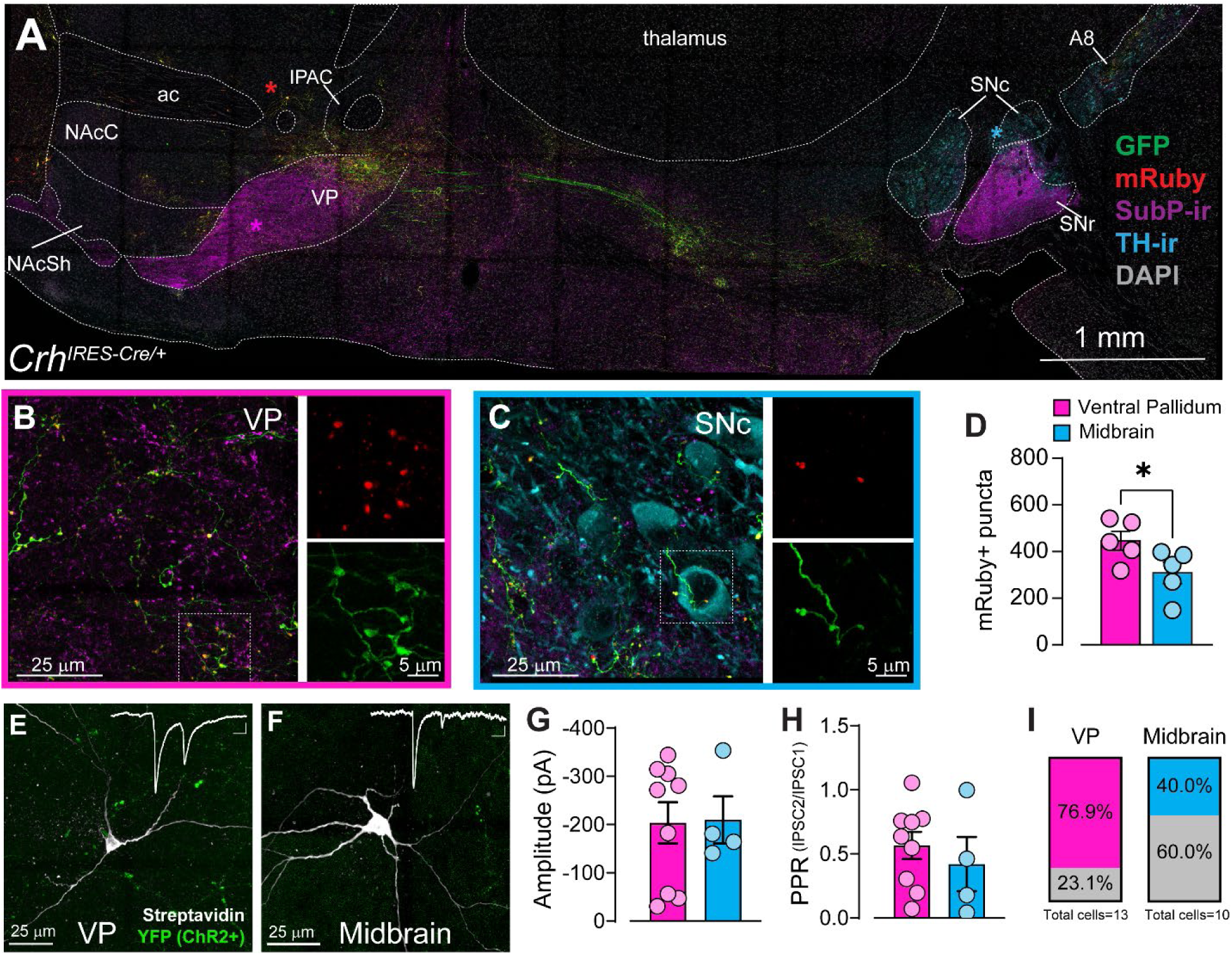
NAc CRF neurons project to ventral pallidal and midbrain regions. (A-C) Representative image of a sagittal section from a *Crh*^IRES-Cre/-^ mouse injected with AAV5-hSyn-FLEX-GFP-Synaptophysin-mRuby into the NAc. Sections were processed for Substance P (SubP, magenta) and tyrosine hydroxylase (TH, cyan) immunoreactivity. GFP+ (green) fibers can be seen along projections, with mRuby+ (red) terminals in (B) ventral pallidum and (C) midbrain. (D) Quantification of mRuby+ puncta in ventral pallidum and midbrain. Significantly more puncta were found in the ventral pallidum, suggesting this is the main projection target of NAc CRF cells (paired t-test, p = 0.0165; N = 5 animals, 3 sections each). (E-F) Post-hoc image from a *Crh*^IRES- Cre/-^ mouse injected with AAV5-DIO-ChR2-eYFP into NAc, with ex vivo electrophysiology recordings performed in the (E) ventral pallidum or (F) midbrain. Recorded cells stained with streptavidin (white) and visualized surrounded eYFP+ (green) terminals (inset, top right). Example traces of optogenetically-evoked IPSCs (oIPSCs) following paired-pulse stimulation (scale bar: 25pA, 50ms). (G) Amplitude of the first oIPSC from paired-pulse recordings of connected cells (displaying a response to optogenetic stimulation) in the ventral pallidum and midbrain (pink and blue, respectively). No difference in amplitude between groups. (H) Paired-pulse ratio (PPR) of oIPSCs in connected cells shows no difference in PPR between downstream regions. (I) Quantification of connectivity in ventral pallidum and midbrain, defined as the percentage of recorded cells that respond to optogenetic stimulation of NAc ChR2+ terminals. Colored portion = portion of connected neurons; gray portion = portion of unconnected neurons. We observed a trend towards higher connectivity in ventral pallidum compared to midbrain (Chi-squared test, p = 0.0721). (Data are represented as mean ± SEM) (ac: anterior commissure, IPAC: posterior limb of the anterior commissure, NAcC: nucleus accumbens core, NAcSh: nucleus accumbens shell, SNc: substantia nigra *pars compacta*, SNr: substantia nigra *pars reticulata*, VP: ventral pallidum).

To confirm synaptic connectivity of these NAc^CRF^ cells, we injected *Crh*^IRES-Cre/-^ mice into the NAc with a channelrhodopsin-expressing, Cre-dependent viral vector. Electrophysiological recordings were performed in the VP and midbrain, with optogenetic stimulation of terminals used to elicit IPSCs (oIPSCs). oIPSCs were found in a subset of both VP and midbrain cells recorded (Figure 2e,f). There were no differences in the amplitude of the first pulse or paired-pulse ratio (Figure 2g,h). However, there was a trend towards a difference in connectivity between regions, with a greater proportion of VP cells responding to photostimulation compared to midbrain (76.9% and 40% of recorded cells, respectively; p = 0.0721) (Figure 2I). Taken together, this data indicates that NAc^CRF^ neurons primarily project to the ventral pallidum, and to a lesser extent the ventral midbrain.

### Electrophysiological examination of NAc^CRF^ neurons reveals three subtypes

The following measures of active and passive conductance were used to classify NAc^CRF^ neurons: membrane resistance, membrane capacitance, resting membrane potential, action potential duration, rheobase, subthreshold I-V slope, F-I curves and sag ratio. While these properties have been most carefully studied in the dorsal striatum (see work by Tepper lab; Surmeier lab (24–28)), work in the NAc has been generally consistent. We assessed these active and passive properties in NAc^CRF^ neurons using two methods: *Crh*^IRES-Cre/-^; Ai14 reporter mice (Figure S4) and *Crh*^IRES-Cre/-^ mice transduced with AAV5-DIO-eYFP viral particles during adulthood. While there was a difference in the distribution depending on methodology, we found three distinct types of electrophysiological profiles present in NAc^CRF^ neurons (Figure 3a).

**Figure 3.**
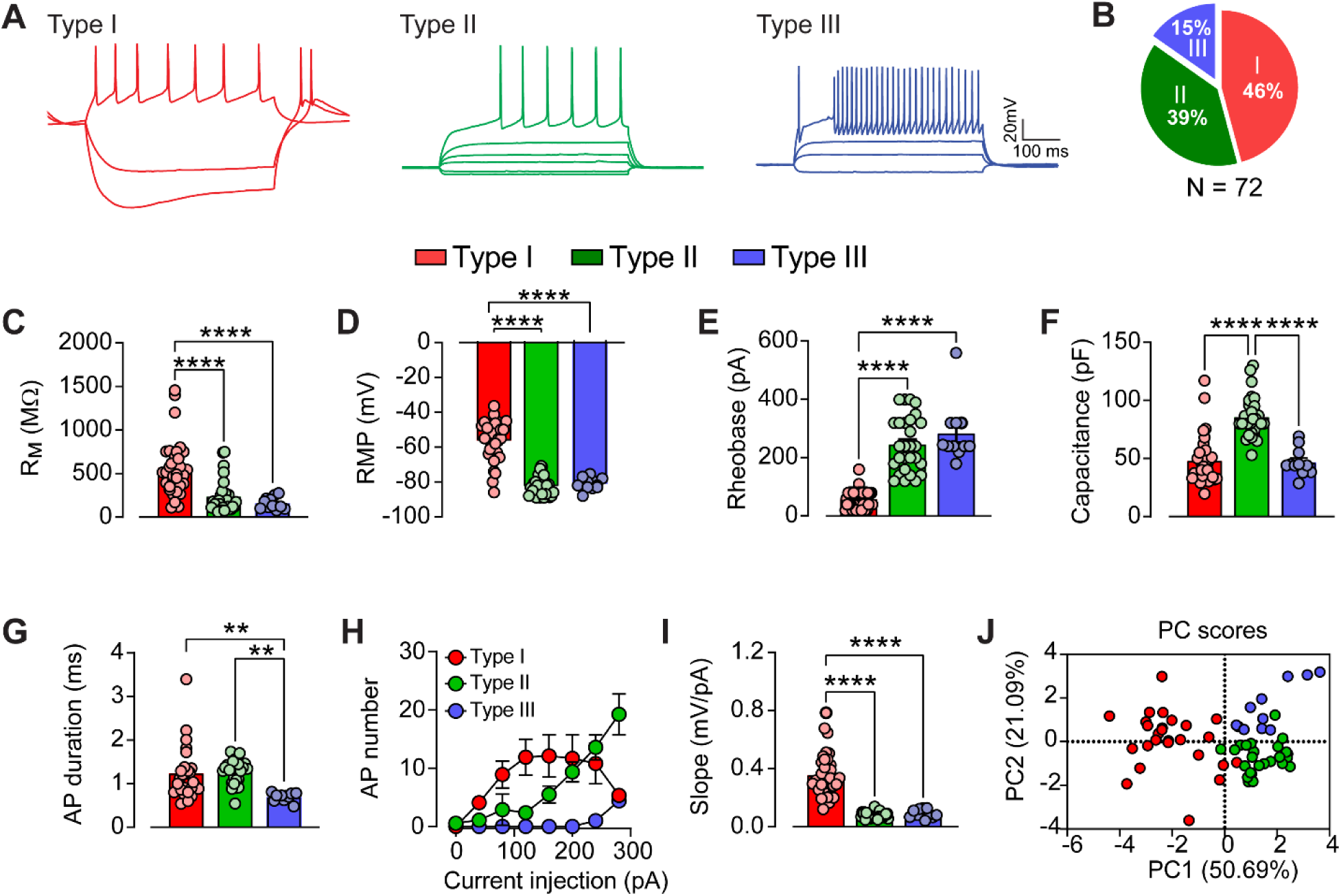
Electrophysiological examination of NAc CRF+ neurons reveals three subtypes. (A) Representative electrophysiological traces of firing patterns of CRF cell subtypes 1, 2 and 3 in response to current injections (-120 to +360, 40pA steps). (B) Proportion of NAc CRF cells that are Type 1 (red; N_cells_ = 33), Type 2 (green; N_cells_ = 28) or Type 3 (blue; N_cells_ = 11). (C) Membrane resistance (R_M_) of CRF cell subtypes. Type 1 cells had significantly higher membrane resistances compared to Type 2 and Type 3 cells. (D) Resting membrane potential (RMP) across CRF cell subtypes. Type 1 cells had significantly higher resting potentials compared to Type 2 and Type 3 cells. Type 2 and Type 3 cells were indistinguishable in this measure. (E) Rheobase of CRF cell subtypes. Type 1 cells had significantly lower rheobases compared to Type 2 and Type 3 cells. (F) Membrane capacitance across CRF cell subtypes. Type 2 cells had significantly higher capacitances compared to Type 1 and Type 3 cells. (G) Average action potential (AP) duration across CRF cell subtypes. Type 3 cells had significantly shorter AP durations compared to Type 1 and Type 2 cells. (H) Firing-current curves (F-I curves) of Type 1, Type 2 and Type 3 cells, showing distinct F-I relationships. (I) Subthreshold IV slope of CRF+ cell subtypes. Type 1 cells had significantly higher slopes than Type 2 and Type 3 cells. (J) Unsupervised principal component analysis of all NAc CRF neurons. Each cell is colored with their experimenter-determined subtype. The three subtypes could be visualized along two PCs, with some overlap between groups. (Data are represented as mean ± SEM) (**p < 0.01, ****p < 0.0001, one-way ANOVAs with Sidak’s).

Type 1, Type 2, and Type 3 NAc^CRF^ neurons displayed differences across a variety of metrics. Across all CRF+ cells analyzed, Type 1 and 2 neurons were more abundant than Type 3 (Figure 3b). Type 1 cells had a greater membrane resistance (one-way ANOVA, F_2,69_ = 18.06, p < 0.0001) (Figure 3c), a more depolarized resting membrane potential (F_2,69_ = 82.64, p < 0.0001) (Figure 3d), as well as a greater variance in resting membrane potentials (Bartlett’s test, p < 0.0001) and a lower rheobase (one-way ANOVA, F_2,69_ = 60.74, p < 0.0001) (Figure 3e) compared to Type 2 or 3. Type 2 neurons had significantly larger membrane capacitance values than Type 1 or 3 (F_2,69_ = 35.54, p < 0.0001) (Figure 3f). Type 3 neurons had a significantly shorter action potential duration compared to Type 1 or 2 neurons (F_2,60_ = 7.433, p = 0.0013) (Figure 3g).

In addition, there was a marked difference in the firing x current (F-I curve) relationship between Type 1, 2 and 3 cells (Figure 3h). Type 1 neurons consistently demonstrated depolarization-induced block of action potential firing at larger depolarizing current injections, whereas Type 2 and 3 neurons did not. Type 1 neurons also had a greater subthreshold IV slope (passive conductance) compared to Type 2 and 3 neurons (F_2,69_ = 60.74, p < 0.0001) (Figure 3i).

To further evaluate experimenter-guided classification of neurons, we performed dimension reduction on the eight electrophysiology parameters using principal component analysis, finding that two principal components (PCs) explained 78.71% of the variance in the data. Type I and Type 2 cells were differentiated mainly along PC1, while PC2 differentiated Type 2 and Type 3 (Figure 3j). However, a lack of clear separation between the different types indicated a degree of overlap. Unsupervised hierarchical clustering analysis revealed that several Type 1 and Type 3 neurons were clustered with Type 2, indicating that a classification approach based on dorsal striatal electrophysiological profiles may not be entirely applicable to the NAc and, in particular, to this CRF population (Figure S5).

In a separate set of experiments, we included biocytin in our recording solution to examine the morphological properties of NAc^CRF^ neurons using a Sholl analysis (Figure S6). Type 1 neurons had significantly fewer intersections than Type 2 or 3 neurons. Likewise, Type 1 neurons had a smaller cumulative dendritic length compared to Type 2 and 3 neurons. There was no significant difference in soma size. Finally, Type 2 neurons had greater spine densities than Type 1 or 3.

To determine whether Type 1, Type 2 and Type 3 neurons are, indeed, projection neurons, we transduced *Crh^IRES-Cre/-^* mice with rgAAV-hSyn-DIO-mCherry or rgAAV-hSyn-DIO-eGFP viral particles into the VP and recorded from fluorescently labeled neurons in the NAc. We found that the electrophysiological phenotypes of these VP-projecting putative CRF neurons reflected that of the larger NAc^CRF^ population, indicating that all three subtypes have the capacity to project out of the NAc (Figure S7).

### NAc^CRF^ SPN subpopulation is more excitable than the totality of NAc SPNs

We sought to offer a possible explanation as to how this population of NAc^CRF^ neurons contained a group of highly excitable, aspiny neurons resembling low-threshold spiking (LTS) interneurons while at the same time not displaying any markers for LTS interneurons (*Npy, Th, Hrt3a, Nos, Sst)* or projection properties that define interneurons. In the dorsal striatum, D1-expressing SPNs recorded during the juvenile period are markedly more excitable, have fewer spines and reduced dendritic length(29). We hypothesized that Type 1 neurons are D1-SPNs that retain their immature excitable profile into adulthood. We compared several key electrophysiological components of D1+ neurons recorded from juvenile (P12–20) *Drd1*^tdTomato^ mice to adult Type 1 and 2 CRF neurons (Figure 4a). Juvenile D1+ neurons exhibited an intermediate phenotype resembling individual properties of either adult CRF Type 1 and 2 neurons.

**Figure 4.**
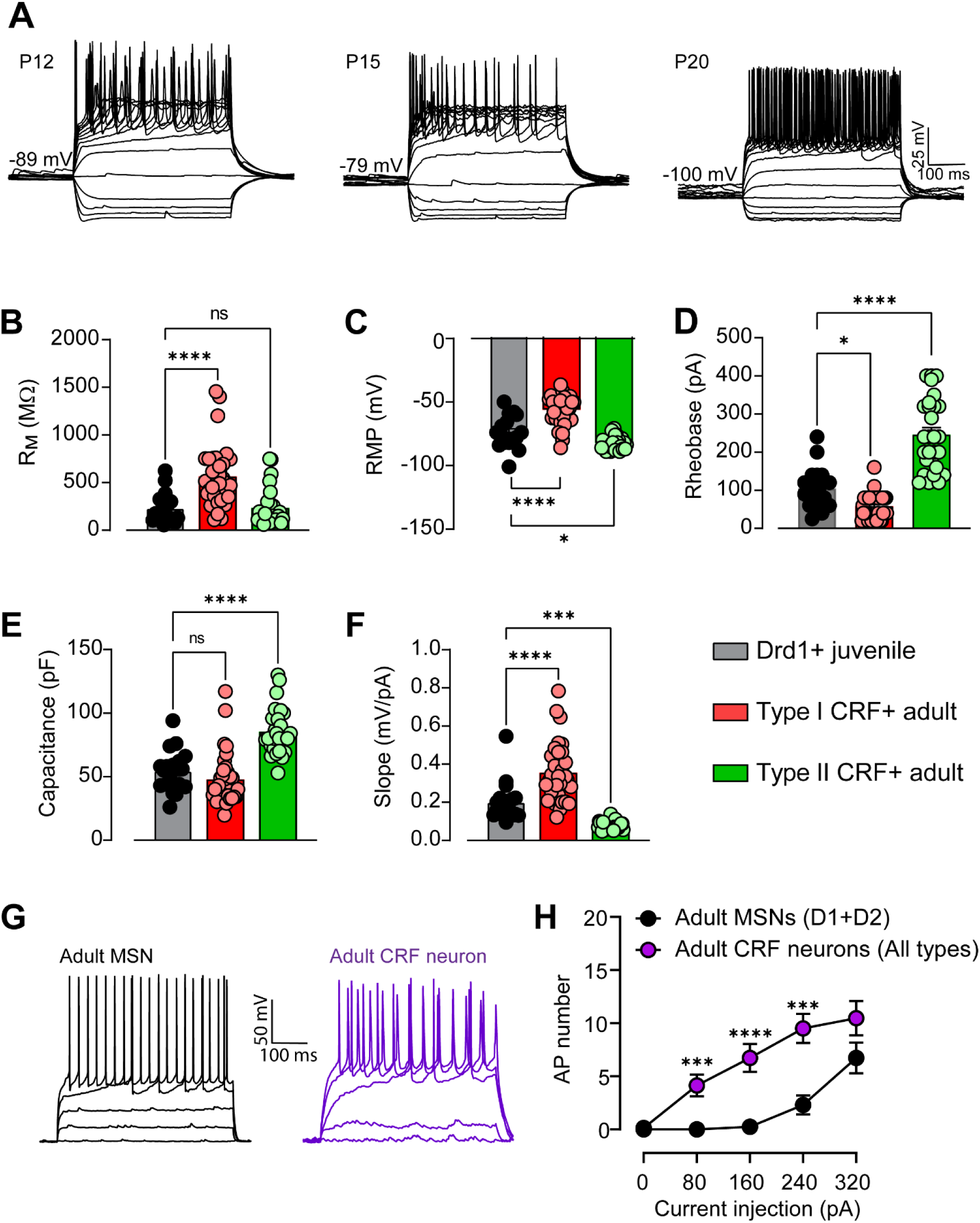
NAc^CRF^ SPN subpopulation is more excitable than the totality of NAc SPNs. (A) Example traces from Drd1+ neurons recorded *Drd1*^tdTomato^ mice of firing patterns in response to a step-current protocol at postnatal days 12, 15 and 20. (B) Membrane resistance in juvenile Drd1+ cells (gray; N_cells_ = 19) and adult CRF+ Type 1 (red; N_cells_ = 33) or Type 2 (green; N_cells_ = 28) cells. Juvenile Drd1+ cells had significantly lower membrane resistances than adult Type 1 cells, but there was no difference between juvenile Drd1+ cells and adult Type 2 cells. (C) Resting membrane potential (RMP) in juvenile Drd1+ cells, adult Type I CRF+ cells, and adult Type 2 CRF+ cells. Juvenile Drd1+ cells had an intermediate phenotype, with their RMP being significantly lower than adult Type 1 cells and significantly higher than adult Type 2 cells. (D) Rheobase of juvenile Drd1+ cells, adult CRF+ Type 1 and Type 2 cells. Similar to RMP, rheobases for juvenile Drd1+ cells was between that of adult Type 1 and Type 2 CRF+ populations. The juvenile Drd1+ rheobase was significantly higher than that of Type 1 cells, but lower than adult Type 2 cells. (E) Capacitances of juvenile Drd1+, adult Type 1 CRF+, and adult Type 2 CRF+ cells. The juvenile Drd1+ cell capacitance was significantly lower than adult Type 2 cells, but not significantly different from that of adult Type 1 cells. (F) The subthreshold IV slope from juvenile Drd1+, adult Type 1 CRF+, and adult Type 2 CRF+ cells. Juvenile Drd1+ cells had an intermediate phenotype, with a slope significantly lower than adult Type 1 cells but significantly higher than the slope of adult Type 2 cells. (G) Example traces of adult SPN neurons recorded from *Drd1^tdTomato^* mouse (gray, left; N_cells_ = 32) and adult NAc^CRF^ neurons recorded from *Crh*^IRES-Cre/-^;Ai14 mouse (purple, right; N_cells_ = 72). (H) Mean firing-current curves (F-I curves) of adult SPNs and adult NAc^CRF^ neurons. CRF+ cells were significantly more excitable compared to SPNs as a whole (mixed effects model with Sidak’s post-hoc). (Data are represented as mean ± SEM) (^ns^p > 0.05, *p < 0.05, ***p < 0.001, **** p < 0.0001, one-way ANOVAs with Dunnett’s post-hoc t-tests unless otherwise noted)

Juvenile D1+ neurons had an average membrane resistance most similar to adult Type 2 CRF neurons, with significantly lower membrane resistances than adult Type 1 cells (one-way ANOVA with Dunnett’s post-hoc t-test; F_2,76_ = 17.33, p < 0.05) (Figure 4b). Juvenile D1+ neurons displayed an intermediate phenotype in both resting membrane potential (RMP) (F_2,77_ = 54.29, p < 0.0001) and rheobase (F_2,78_ = 66.66, p < 0.0001). The RMP of juvenile D1+ neurons was significantly lower than that of adult Type 1 CRF neurons, but was significantly higher than the RMP of adult Type 2 CRF neurons (Figure 4c). The rheobase of juvenile D1+ neurons was significantly higher than that of adult Type 1 and significantly lower than adult Type 2 CRF cells (Figure 4d). While the membrane resistance of juvenile D1+ neurons more closely resembled that of adult Type 2 CRF neurons, the membrane capacitance of juvenile D1+ neurons was more like the capacitance of adult Type 1 CRF neurons. The membrane capacitance of juvenile D1+ neurons was significantly lower than that of adult Type 2 neurons, but not significantly different from Type 1 neurons (F_2,77_ = 32.77, p < 0.0001) (Figure 4e). These findings are consistent with the notion that there are fewer spines and therefore less overall surface area in immature D1+ neurons compared to fully mature SPNs. Finally, the subthreshold IV slope of juvenile D1+ cells was significantly different from that of both adult CRF neuron groups, displaying an intermediate phenotype. Adult Type 1 CRF neurons had a significantly higher slope than juvenile D1+ neurons, while adult Type 2 CRF neurons had a significantly lower slope than juvenile D1+ neurons (F_2,77_ = 48.37, p < 0.0001) (Figure 4f).

We conclude that CRF+ cells are SPNs with both mature and immature electrophysiological profiles. To investigate how CRF neuronal excitability compared to NAc SPN average excitability, we recorded from adult *Drd1*^tdTomato^ mice in which we acquired 16 D1+ and 16 D1-(putative D2+(18)) SPNs. NAc^CRF^ putative SPNs as a population were significantly more excitable than randomly sampled SPNs (mixed effects model, fixed effects Type III, type x time interaction, F_4,379_ = 4.419, p = 0.0017, Sidak’s post-hoc t-tests, p < 0.001) (Figure 4g,h). This further supports the conclusion that the CRF+ population retains their excitable, juvenile phenotype at a higher rate than SPNs as a whole.

### NAc^CRF^ neurons track reward outcomes in an operant learning task

Optogenetic stimulation of NAc^CRF^ neurons is sufficient to support intracranial self-stimulation and promote the incentive salience of sucrose reward(12). However, the activity patterns of the NAc^CRF^ neuronal population during effortful reward learning has not been studied. To do this, we transduced AAV5-FLEX-GCaMP6s into the NAc of *Crh*^IRES-Cre/-^ mice and implanted optical fibers unilaterally into the same location (Figure 5a;S8). We recorded NAc^CRF^ neuronal activity patterns every other day during an instrumental conditioning task (Figure 5b). On average, mice took 5 days to meet criteria in both FR1 and FR3 (Figure 5c). There was a significant difference in the fluorescence elicited by rewarded verses unrewarded head entries (two-way ANOVA with Sidak post-hoc; main effect of time F_13,19_ = 18.12, p < 0.001; group F_1,7_ = 5.362, p = 0.0537; time x group F_13,91_ = 5.488, p < 0.0001) (Figure 5d,e). Rewarded head entries elicited significantly more activity immediately following magazine entry (0 to 2.5 s from TTL) compared to unrewarded entries. Interestingly, this increased activity in response to rewarded head entries only was followed by a significantly greater depression in activity which persisted for several seconds (5 to 15 s from TTL).

**Figure 5.**
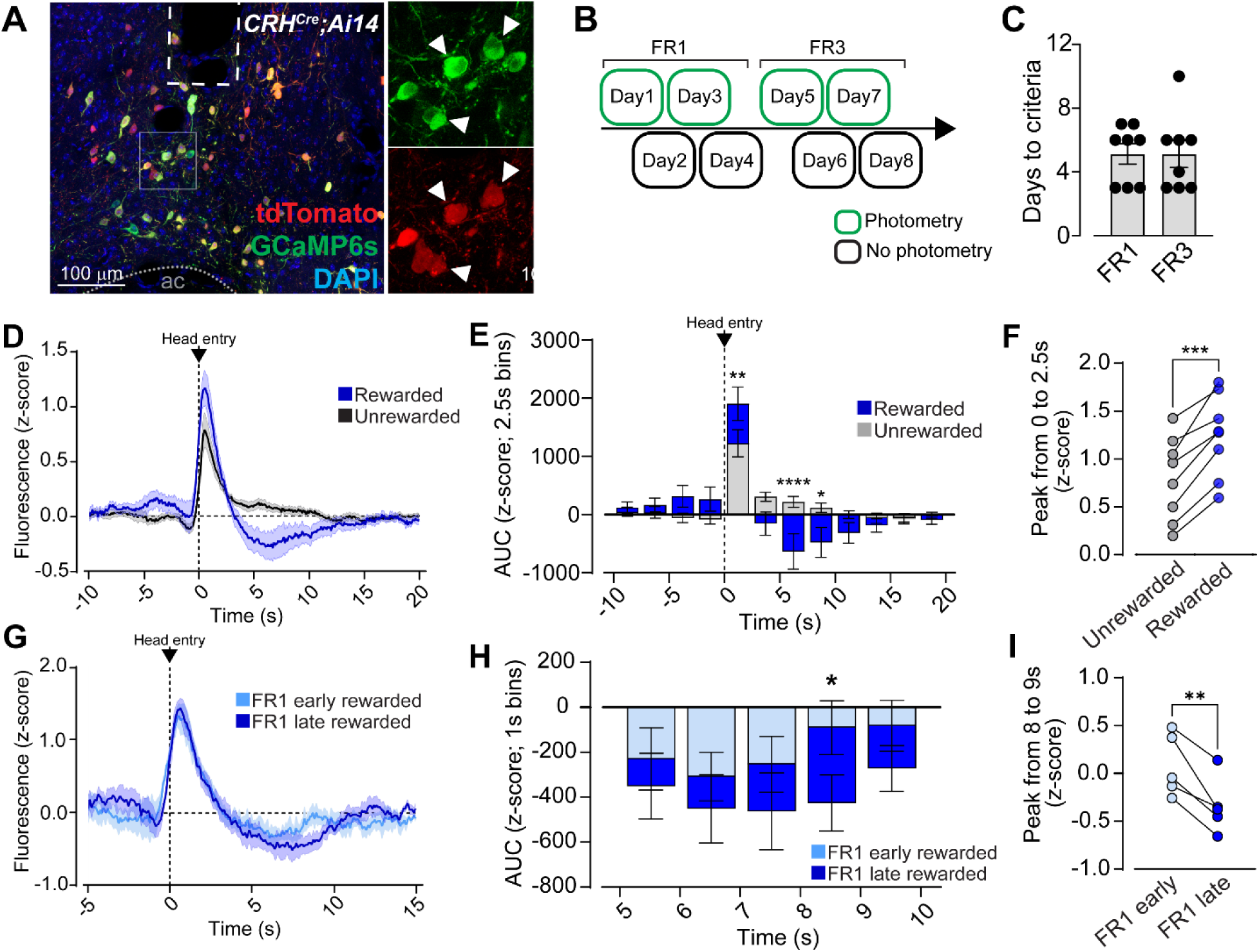
NAc^CRF^ neurons track reward outcomes in an operant learning task. (A) Representative image of GCaMP6s (green) expression below the fiber optic tip (dotted line) in the NAc of *Crh*^IRES-Cre/-^;Ai14 mice injected with AAV5-Syn-FLEX-GCaMP6s. Endogenous Cre-driven tdTomato expression (red) can be seen co-localized with GCaMP6s viral particles (green), indicating recordings were of putative CRF+ cells (inset, white arrows). Cells were counterstained with DAPI (blue). (B) Schematic of operant behavior task training and recording schedule. Mice were trained to a fixed ratio 1 (FR1) criteria, then fixed ratio 3 (FR3) criteria. Fiber photometry recordings were performed every other day. (C) Number of training days required for mice to reach criteria on FR1 and FR3 schedules. (D) Average fluorescence (z-score) from NAc CRF cells time-locked to head entry into the reward magazine (t=0). Head entries were separated into those immediately following reward delivery when the reward was present in the magazine (rewarded; blue) and head entries without reward delivery (unrewarded; black) (N_mice_ = 8). (E) Area under the curve (AUC) for GCaMP6s fluorescence z-scores binned into 2.5 s increments, time-locked to head entry into reward magazine. Rewarded head entries (blue) were accompanied by significantly higher fluorescence than unrewarded entries (gray) immediately following magazine entry, followed by significantly lower fluorescence 5–10 s after magazine entry (two-way ANOVA). (F) Within-subject comparison of the average peak fluorescence immediately following magazine head entry (0 to 2.5 s after entry) for each mouse. Rewarded head entries elicited significantly greater peak calcium responses compared to unrewarded head entries for every mouse (paired t-test). (G) Average fluorescence (z-score) time-locked to rewarded head entry into magazine (t=0) for early (days 1-2; light blue) and late (days 3-4; dark blue) FR1 training days (N_mice_ = 4). (H) AUC for fluorescence z-score binned into 1 s increments for rewarded head entries during early (light blue) and late (dark blue) FR1 training days during the post-retrieval inhibition period (5 to 10 s following head entry TTL). There is a significant difference between early and late training phases, with increased inhibition during late FR1 training (two-way ANOVA). (I) Within-subject comparison of the average peak fluorescence during early and late FR1 training during latter half of inhibition period (8 to 9 s) following rewarded head entry TTL. Peak fluorescence during this period is suppressed in late FR1 training compared to early FR1 training (paired t-test). (Data are represented as mean ± SEM) (*p < 0.05, **p < 0.01, ***p < 0.001, **** p < 0.0001)

We examined the development of this activity pattern during early training (days 1-2 compared to days 3-4 during FR1). Peak responses to rewarded head entries were not different across training. However, there was a significant difference in post-retrieval inhibition between early and late FR1 training days (main effect of time F_4,16_ = 4.755, p = 0.0101; day F_1,4_ = 9.038, p = 0.0397; time x day F_4,16_ = 0.6189, p = 0.6554), with increased inhibition in later FR1 days verses early FR1 days (Figure 5g-i).

### CRF release from NAc^CRF^ neurons constrains action-outcome acquisition

We next wanted to determine the role of CRF peptide release from NAc neurons during this same operant task. We used two different methods. First, *Crh*^loxP/loxP^ mice were transduced with AAV5-CMV-Cre-eGFP (CRF cKO) bilaterally into the NAc to conditionally knock out CRF peptide from NAc neurons. AAV5-CMV-eGFP was used as the control. We validated the knock-down using RNAscope and qPCR, and confirmed this manipulation did not alter hypothalamic-pituitary-adrenal axis function or body weight (Figure 6a,b;S9).

**Figure 6.**
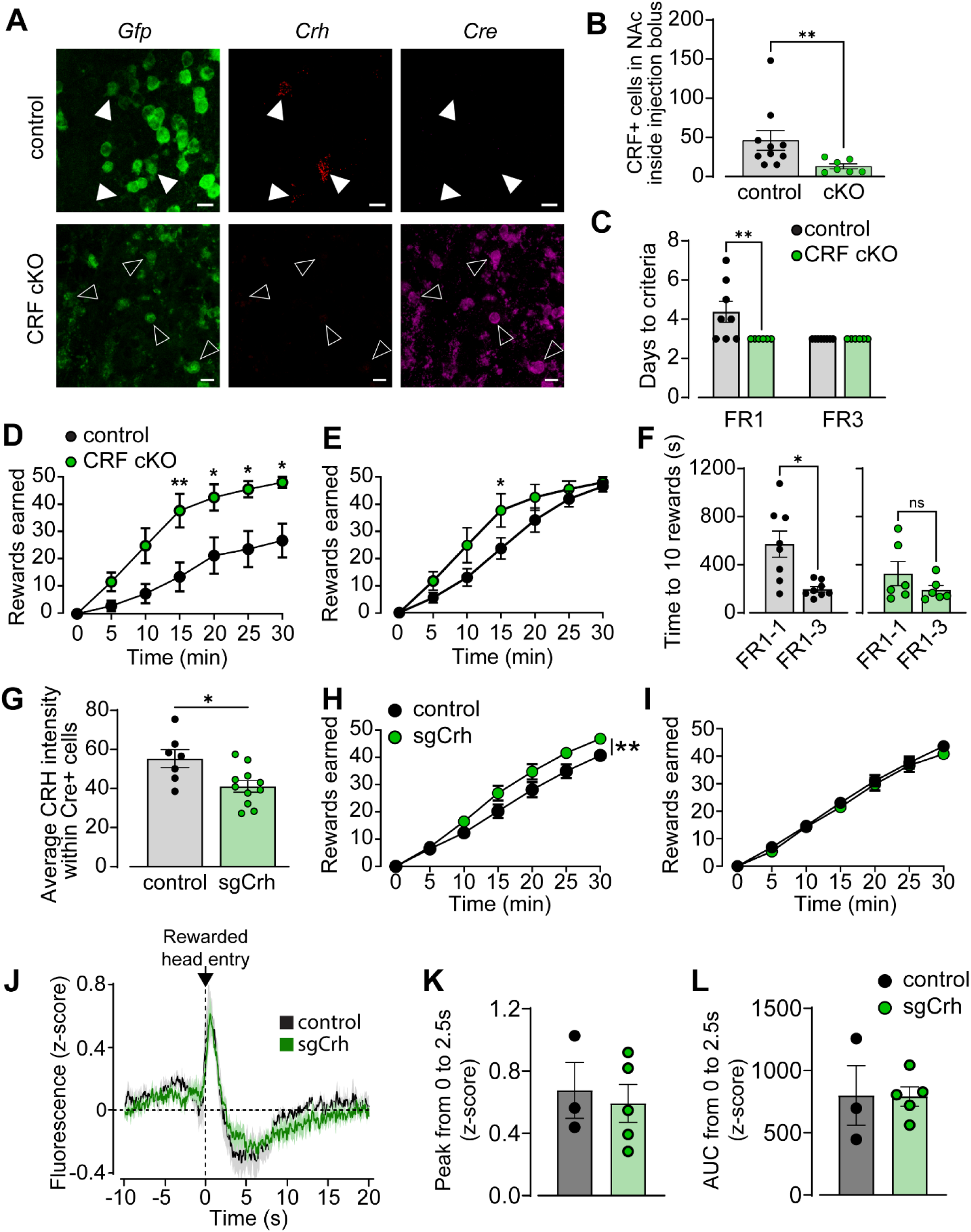
CRF release from NAc^CRF^ neurons constrains action-outcome acquisition. (A-F) Conditional knockout (cKO) of NAc^CRF^ was achieved by bilateral injection of AAV8-CMV-eGFP (control) or AAV8-CMV-eGFP-Cre (CRF cKO) virus into the NAc of *Crh*^loxP/loxP^ mice. (A) Representative images from control (top) and CRF conditional knockout (bottom) mice. Injections were validated by in situ hybridization for *Gfp* (green), *Crh* (red), and *Cre* (magenta) mRNA. In control mice, there was co-localization between *Gfp* and *Crh* (closed arrows), while CRF cKO animals had co-localization between *Gfp* and *Cre* (open arrows) but diminished co-localized of *Crh*. (B) Quantification of NAc CRF+ cells within the injection bolus of control (gray) or CRF cKO (green) mice. There was a significant reduction in the number of *Crh*+ cells in cKO mice (Mann-Whitney). (C) Days of testing required to reach criteria in an operant task with FR1 or FR3 reward schedule in control and CRF cKO mice. CRF cKO mice were significantly quicker to reach criteria in FR1 training. (D) Number of rewards earned over time on the first unbaited day of FR1 training and (E) on the first FR1 day in which criteria was reached. CRF cKO mice were significantly quicker to acquire rewards compared to controls (N_mice_ = 6 and 8, respectively). (F) Average time to earn 10 rewards in FR1 training on the first day (FR1-1) and third day (FR1-3) daily criteria was met. Control mice (gray, left) significantly reduced their time to 10 rewards across training, earning rewards faster on the third day. CRF cKO mice (green, right) did not show a difference in time to acquire 10 rewards between FR1 training days (paired t-test). (G-L) *Crh*^IRES-Cre/-^ mice were injected bilaterally into NAc with either control (AAV5-FLEX-SaCas9-U6-sg*ROSA26*) or sgCrh (AAV5-FLEX-SaCas9-U6-sg*Crh*) virus, along with GCamp6s virus (AAV5-Syn-FLEx-GCaMP6s-WPRE) and implanted with an optic fiber above the injection site. Mice were trained in a fixed ratio 1 (FR1) task, then a fixed ratio 3 (FR3) task, and finally a 5-day reversal test. Fiber photometry recordings were performed every other day. (G) CRISPR-mediated knockdown of CRF was validated using *in situ* hybridization in a cohort of mice injected with either sgCrh or control virus. Average intensity of *Crh* mRNA within Cre+ cells (putative CRF+) was quantified. sgCrh injection significantly reduced the average intensity of *Crh* in Cre+ cells compared to control (t-test). (H) Average performance during FR1 task days, measured as rewards earned over session time. Data averaged by mouse, including only the four consecutive days the animal met FR1 criteria immediately prior to moving to FR3 task. sgCrh mice (green; N = 6) earned rewards significantly quicker than controls (black; N = 5). (I) Average performance during FR3 task, including only the four consecutive days mice met criteria before being moved to reversal test. There was no difference between control and sgCrh groups speed to acquire rewards in the FR3 task. (J) Fiber photometry recordings from FR1 and FR3 days at criteria were averaged by mouse for control (gray; N = 3) and sgCrh (green; N = 5) groups. Average fluorescence (z-score) from NAc CRF cells time-locked to head entry into the reward magazine when a reward was present in the magazine (rewarded head entry) for control and sgCrh mice during fixed ratio (FR1 and FR3) task. (K) Peak fluorescence and (L) AUC from 0 to 2.5 s following rewarded head entry did not differ between control (black) and sgCrh (green) mice (t-tests). There were no differences between groups in activity response to rewarded head entries. (Data are represented as mean ± SEM. Scale bar = 10 µm) (^ns^p > 0.05, *p < 0.05, **p < 0.01, two-way repeated measures ANOVA with Sidak’s post-hoc unless otherwise noted above.)

CRF deletion from NAc neurons in adulthood resulted in a faster time to reach criteria for FR1, which did not persist to FR3 training (two-way repeated measures ANOVA with Sidak’s; main effect of FR schedule: F_1,12_ = 4.900, p = 0.047; group: F_1,12_ = 4.900, p = 0.0470; and FR schedule x group: F_1,12_ = 4.900, p = 0.0470) (Figure 6c). During the first unbaited FR1 training day, CRF cKO mice had quicker reward acquisition (two-way repeated measures ANOVA with Sidak’s; main effect of time: F_6,72_ = 51.21, p < 0.0001; group: F_1,12_ = 8.242, p = 0.0141; time x group: F_6,72_ = 4.804, p = 0.0004) (Figure 6d). We also compared reward acquisition over session time on the first day criteria was reached, and found that CRF cKO mice were quicker to acquire rewards at the beginning of training even when all mice were reaching criteria (two-way repeated measures ANOVA with Sidak’s; main effect of time: F_6,72_ = 140.0, p < 0.0001; and time x group: F_6,72_ = 2.933, p = 0.0129) (Figure 6e). This effect did not persist across subsequent days, with no difference in overall nose pokes between groups on any day (Figure S10). In line with our data showing an initial difference in speed of reward acquisition, control mice significantly reduced their time to earn 10 rewards between the first day criteria was reached on FR1 and the third day (paired t test, p = 0.014). There were no differences in reward earning during FR3 or a progressive ratio task (Figure 6c; S10). Thus, deletion of CRF from the NAc appears to promote quicker initial acquisition of operant reward learning. This effect is not due to gross changes in reward preference since control and CRF cKO mice showed no differences in the sucrose preference test (Figure S10). We did, however, find that CRF cKO mice have a reduced psychomotor response to novelty in an open field, indicating that, normally, CRF release from NAc^CRF^ neurons promotes exploration (Figure S10).

Another consideration is that CRF deletion may produce long-term changes in NAc^CRF^ neuronal function in addition to reduced CRF release itself, that lead to deficits in reward learning. To test this, we utilized a Cre-dependent CRISPR virus directed at *Crh* (sg*Crh*) and accompanying control virus (sg*Rosa*) to knockdown NAc^CRF^. The benefit of this method is that it allows for co-expression of fluorophores and GECIs in cells receiving CRISPR virus. CRF knock-down was validated using fluorescent in situ hybridization (Figure 6g). Using ex vivo electrophysiological techniques, we confirmed that excitability, glutamatergic synaptic transmission and GABAergic synaptic transmission were not significantly changed following CRF knock-down (Figure S11). As seen in the CRF cKO experiments, sg*Crh*-mediated knock-down increased the speed of reward acquisition in FR1 (two-way repeated measures ANOVA with Sidak’s; main effect of time: F_6,54_ = 393.6, p < 0.0001; time x group: F_6,54_ = 3.171, p = 0.0097), with no difference in acquisition on FR3 days (Figure 6h,i). Importantly, CRF knock-down did not produce significant changes in NAc^CRF^ neuronal activity patterns during this task (Figure 6j-l;S12).

### CRF deletion produces deficits in reversal learning

We performed a 5-day reversal experiment to determine if behavioral flexibility differed between groups. Compared to controls, sg*Crh* mice were significantly slower to earn rewards in the reversal sessions on average (two-way repeated measures ANOVA with Sidak’s; main effect of time: F_6,36_ = 233.3, p < 0.0001; group: F_1,6_ = 6.163, p = 0.0476; time x group: F_6,36_ = 2.593, p = 0.0342) (Figure 7a). Importantly, this difference persisted across the reversal trials, with sg*Crh* mice being significantly slower to earn rewards on both the initial (main effect of time: F_6,36_ = 42.64, p < 0.0001; group: F_1,6_ = 8.620, p = 0.0261; time x group: F_6,36_ = 3.582, p = 0.0069) (Figure 7b) and final (main effect of time: F_6,36_ = 128.3, p < 0.0001; group: F_1,6_ = 8.289, p = 0.0281; time x group: F_6,36_ = 4.395, p = 0.0020) (Figure 7c) reversal days, and there was no difference in the number of pokes to the inactive (previously active) port (main effect of time: F_6,36_ = 27.48, p < 0.0001; group: F_1,6_ = 0.0028, p = 0.9592; time x group: F_6,36_ = 0.4647, p = 0.8297) (Figure 7d). Together, these data indicate a deficit in updating reward contingencies.

**Figure 7.**
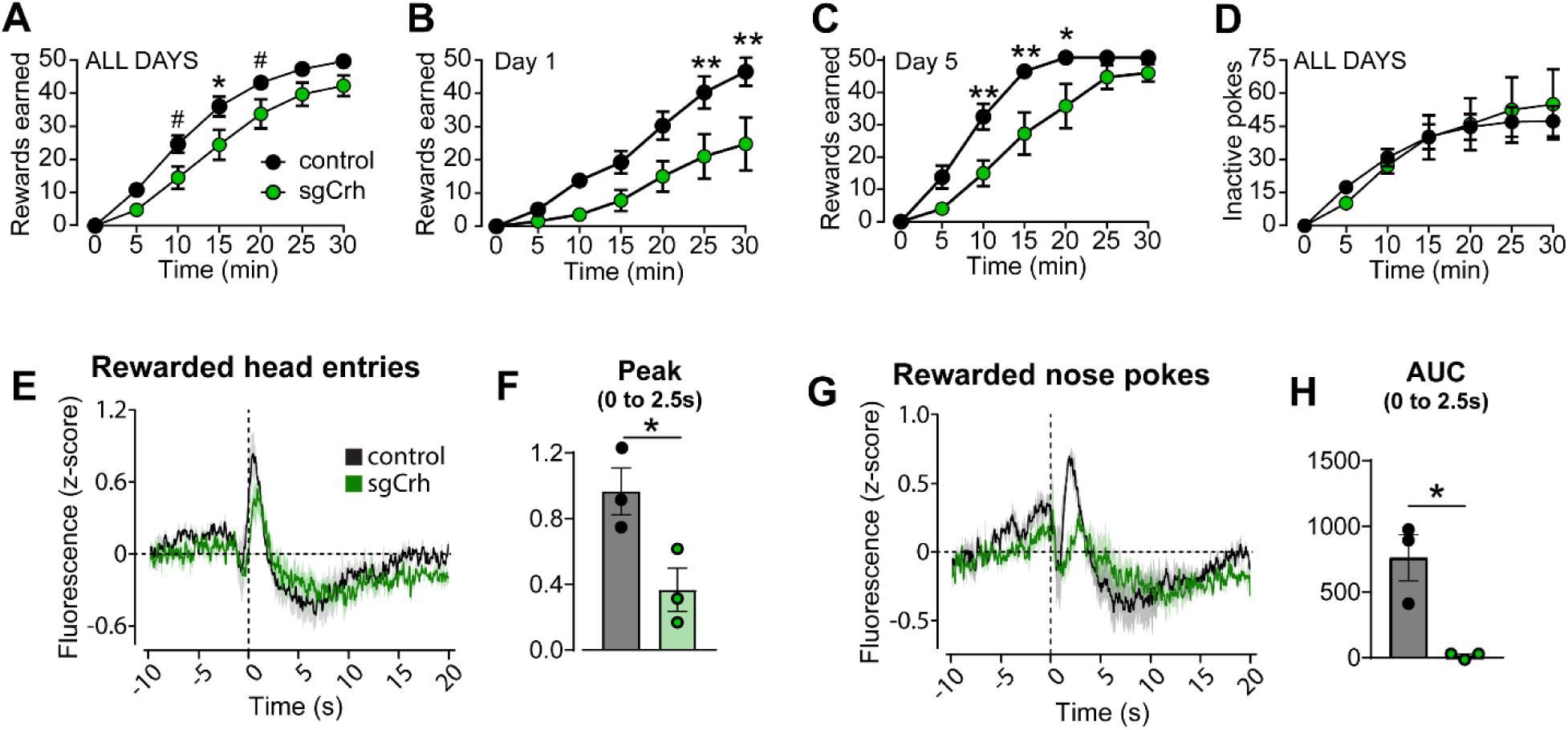
CRF deletion produces deficits in reversal learning. Following successful FR3 training, control (black) and sgCrh (green) mice were put through 5 days of reversal testing wherein the active and inactive nose ports were switched. (A) Rewards earned over session time during reversal test, averaged by mouse across all reversal days. sgCrh mice were significantly slower to earn rewards compared to controls (N = 4 and 4). (B) Rewards earned over session time during the first reversal day. sgCrh mice were significantly slower to earn rewards on the first reversal day. (C) Rewards earned over session time during the final reversal day. As during the first day, sgCrh mice were significantly slower to earn rewards in the final reversal session. (D) Average inactive pokes over session time during reversal test days, averaged by mouse for all reversal days. There was no difference between groups in inactive pokes. (E-H) Fiber photometry recordings from reversal test days were averaged by mouse for control (gray) and sgCrh (green) groups. Peak and area under the curve (AUC) measures were taken from 0 to 2.5s following behavioral event TTL (TTL at t=0). (E) Average fluorescence (z-score) from NAc CRF cells in response to rewarded head entry for control and sgCrh mice during reversal test. (F) Peak of fluorescence following rewarded head entry. In reversal sessions, NAc CRF cells from control mice had a significantly higher peak compared to controls in response to rewarded head entries (t-test). (G) Average fluorescence (z-score) following a rewarded nose poke in reversal test sessions for control and sgCrh mice. (H) AUC of fluorescence following a rewarded nose poke. AUC of fluorescence following a rewarded nose poke is significantly higher in control mice compared to sgCrh mice in reversal sessions (t-test). (Data are represented as mean ± SEM.) (^#^p < 0.1, *p < 0.05, **p < 0.01, two-way repeated measures ANOVA with Sidak’s post-hoc unless otherwise noted above.)

We observed several differences in NAc^CRF^ activity patterns in mice with CRF knock-down compared to control during reversal learning. The amplitude of the peak to rewarded head entries was significantly diminished in sg*Crh* mice (t-test, p = 0.0368), but there were no significant differences in the AUC (Figure 7e,f;S12). Unrewarded head entries did not differ significantly between groups (Figure S12). However, contrary to the fixed ratio sessions, the pattern of activity following a rewarded nose poke in reversal sessions was significantly different (Figure 7g). While the calcium peak was not significantly different, the AUC of activity 0 to 2.5 s following a rewarded poke was robustly diminished in sg*Crh* mice compared to controls (t-test, p = 0.0134) (Figure 7h). Taken together, these data suggest that the lack of NAc^CRF^ neuronal functionality may alter the response of these cells to the acquisition of reward in this reversal task. Thus, we cannot rule out the possibility that deficits in reversal learning are due to both a reduction in NAc^CRF^ function overall and reduction in CRF release.

## Discussion

In this study, we comprehensively analyzed the transcriptomic, electrophysiological and morphological features of NAc^CRF^ neurons and provide evidence that this population constrains action-outcome acquisition during reward learning by promoting environmental exploration. To our knowledge, the neurochemical and physiological identity of these neurons has never been determined, nor has the behavioral role of CRF release itself from these neurons. NAc^CRF^ neurons are a subpopulation of SPNs that mostly contain D1 and to a lesser extent D2 receptors, are enriched in “eccentric” SPN markers, primarily project to the VP, and are more excitable than SPNs as a whole. NAc^CRF^ neurons increase their activity during both unrewarded and rewarded head entries, but the integrated calcium response was greater for rewarded head entries. Furthermore, activity is significantly depressed only following rewarded head entries, and this depression evolves over training over a similar timescale to the diminishment of the effects of CRF deletion. Conditional CRF deletion enhanced the acquisition of the instrumental task and increased the number of rewards acquired during the initial acquisition phase, an effect that was consistent across two knock-down methodologies. These findings indicate that CRF release from this sparse population of NAc neurons constrains reward acquisition during a period when action-outcome patterns are being established. In addition, there is decreased reward earning in a reversal task following CRF knockdown This suggests that CRF may play a role in preventing hyperfocus (or encouraging flexibility) during certain behaviors that allows for exploration of the environment.

### CRF neurons in the NAc are comprised of a heterogenous population of SPNs

Neurochemically, the majority (70–85%) of CRF cells expressed SPN markers with a small percentage displaying the fast-spiking interneuron neurochemical phenotype. Analysis of projection patterns supported the neurochemical data. Despite being heavily comprised of D1-only expressing SPNs, the VP projection is more robust than the projection to midbrain. These data advance our understanding of NAc^CRF^ neurons, as it has been presumed that these neurons produce their behavioral output through local connections. Since SPNs send projections outside the NAc while also maintaining a dense lateral plexus via axon collaterals, they can influence both local circuit function and long-range projection regions.

Our electrophysiological and morphological characterization of NAc^CRF^ neurons challenges the current striatal cell classification system that was developed primarily in the dorsal striatum. Approximately half of the putative NAc^CRF^ cells in our preparation displayed a non-canonical SPN electrophysiological and morphological phenotype more characteristic of low-threshold spiking interneurons (i.e. NPY-, SST-, 5-HT3- and TH-containing interneurons)(24–26). This was in direct contrast to our neurochemical examination, which found negligible expression of these markers in NAc^CRF^ neurons. A heterogeneous composition of CRF neurons is consistent with observations in both the BNST and hippocampus (30, 31). Hippocampal CRF neurons are comprised of several different interneuron subtypes (e.g. PV+, CCK+, CR+)(31). Likewise, CRF neurons in the BNST have been clustered into three different types based on their electrophysiological properties(30). It is therefore not surprising that NAc^CRF^ neurons do not “follow the rules”. Further, a recent study demonstrated that gastrin-releasing NAc neurons were SPNs with a similar non-canonical excitable profile (32). Importantly, our retrograde viral experiments combined with electrophysiology demonstrate that all three electrophysiological subtypes can project to VP, again supporting the notion that these are primarily SPNs not interneurons. We offer the following as a possible mechanism for the heterogenous composition. D1-SPNs in the dorsal striatum are more excitable and aspiny prior to adulthood(29). We find that Type 1 and Type 2 CRF neurons recorded from adult mice share electrophysiological features with D1-containing direct SPNs recorded from juveniles. Some portion of these D1-SPNs may retain some of their “immature” properties in the NAc (for additional discussion, see *Supplemental Discussion*). A key point of this analysis is that NAc^CRF^ putative SPNs are more excitable than SPNs overall. This suggests a potential role in encoding saliency in environment, a notion we plan to explore in future studies.

### Accumbal CRF neuronal activity tracks and shapes action-outcome during reward learning

This study demonstrates a critical role for CRF in instrumental reward learning, particularly during early learning and changing contingencies, which has not previously been incorporated into models of NAc function and reward. CRF acts to constrain action-outcome learning under relatively normal conditions (i.e. not during severe or repeated stressor exposure). Hedonic perception or motivational drive do not seem to play a role in this context, as CRF cKO mice show normal sucrose preference and show no difference in a progressive ratio task once fully trained. We did find that CRF promotes exploratory behavior in response to a novel open field environment. Further, we find that CRF plays a role in contingency updating a reversal task. We originally hypothesized that deletion of CRF would impair reward learning and action-outcome performance in food restricted animals based on work demonstrating that exogenous CRF application enhances acetylcholine and dopamine transmission in the NAc and facilitates appetitive behaviors(13–15, 17, 33). We have disproven this original hypothesis and instead offer the following interpretation (for additional interpretations see *Supplemental Discussion*).

We posit that CRF release from NAc^CRF^ neurons enhances exploratory behavior thus limiting focus on one specific task, especially early in task acquisition. Exogenous application of CRF into the NAc enhances motor behavior, while inhibition of CRF receptors in the NAc or deletion of CRF receptors from dopamine neurons inhibits exploration of novel objects(13, 34, 35). Here, we find that deletion of CRF from accumbal neurons reduces locomotor responses to a novel environment, consistent with previous findings. Further, our finding that a loss of CRF impairs reversal learning adds support to this flexibility interpretation. The notion that CRF enhances cognitive flexibility by enhancing exploration or environmental surveillance has been suggested in the context of CRF excitation of noradrenergic locus coeruleus neurons that project to the prefrontal cortex(36). A similar role for CRF has been assigned to CRF modulation of serotonin(36). We conclude that CRF regulation of accumbal, pallidal and midbrain function may infuse some flexibility in engaging and disengaging with reward learning that is absent following deletion of CRF from NAc neurons.

## Acknowledgements

This study was funded by K99/R00 Pathway to Independence award (MH109627 JCL), NIMH BRAINS R01 (MH122749 JCL), and NIDA F31 NRSA (DA059436 EAE). This work was supported by NIDA Core “Center of Excellence” awards to the University of Minnesota (P30DA048742) and University of Washington (P30DA048736). This work was supported by the resources and staff at the University of Minnesota University Imaging Centers (UIC, SCR_020997). We thank Kasey Bertelsen for assistance with the spontaneous PSC analysis. We thank Jennifer Robeson for assistance with behavioral analysis. We thank Dr. Veronica Alvarez for the use of equipment and reagents. We thank Mariah Blegen for assistance with qPCR. We thank Rachel Dick for technical assistance. Dr. Kavya Devarakonda contributed key edits to the manuscript. The authors have uploaded a preprint of this manuscript to BioRxiv prior to publication. The authors declare no competing financial interests.

## Author Contributions

conceptualization: E.A.E. and J.C.L.; formal analysis: E.A.E., A.E.I., J.J.S., C.L.P., M.F., J.M.R., A.M.R., E.M.K., J.C.L.; funding acquisition: E.A.E., J.C.L.; investigation: E.A,E, A.E.I, J.J.S, C.L.P, M.F., J.M.R, A.M.R., E.M.K, J.C.L; methodology: E.A.E., A.E.I., C.L.P., J.C.L.; project administration and supervision: E.A.E., A.E.I., C.L.P., J.C.L.; resources: S.S., L.S.Z, M.S.C., J.C.L.; software: E.A.E., A.E.I., J.C.L.; visualization: E.A.E., A.E.I., J.C.L.; writing – original draft: E.A.E., J.C.L.; writing - review & editing: E.A.E., A.E.I., J.C.L.

## Supplemental Materials

## Supplemental Methods

### Animals

All procedures were done in accordance with the University of Minnesota Animal Care and Use Committee. Both male and female mice were used in approximately equal numbers, ages post-natal day 60-224, unless otherwise specified. Mice were group housed in specific-pathogen free conditions under a 16h light cycle (6:00 ON/20:00 OFF) with freely available food and water; “summer” photoperiods to facilitate breeding. Two weeks prior to experimental testing, mice were transferred to investigator managed standard housing and kept under a 12h light cycle (6:00 ON/18:00 OFF) for the remainder of testing. Unless otherwise noted, food and water were available ad libitum. For in situ hybridization experiments, C57BL/6J mice (JAX stock: 000664) were used. For viral tracing experiments, optogenetic and CRISPR experiments, *Crh*^IRES-Cre/-^ mice (JAX stock: 012704) were used. For electrophysiology and fiber photometry experiments, *Crh^IRES-Cre/-^* mice were crossed with Ai14 (JAX stock: 007914) reporter mice. Although this *Crh^IRES-Cre/-^* line has been used extensively, it has not been validated for the NAc. The specificity of this line (defined as the % of *Crh+Cre+*/total *Cre+* cells) was very high (>85%) for the NAc and similar in all regions analyzed: the paraventricular nucleus of the hypothalamus (95 ± 1%), NAc (88 ± 4%), BNST (93 ± 2%) and central amygdala (95 ± 4%) (Figure S1). In contrast, the penetrance of Cre (defined as (%*Crh+Cre*)/total *Crh*) was significantly lower in the NAc (38 ± 4%) compared to the paraventricular nucleus (91 ± 1%), BNST (92 ± 2%) and central amygdala (78 ± 6%) (Figure S1). For CRF conditional knockout experiments, *Crh*^loxP/loxP^ mice generated by Dr. Michael S. Clark and validated by Dr. Larry Zweifel at the University of Washington were used(1).

### Rigor and Reproducibility

*Blinding*: When possible, experimenters directly performing the experiments were blinded to animal conditions: experimental or control.

*Automation*: Wherever possible, we used automated systems to acquire and analyze our data. *Power analyses* were used to determine sample sizes using G*Power 3.1(2, 3). The power analysis is based on the expected magnitudes and SDs of results based on our prior data and previous literature as well as an α-value of 0.05. Of note, for the majority of experiments, we did not fully power our experiments to assess sex differences. In the future we are analyzing potential trends and conducting fully powered sex-difference experiments.

### Fluorescent in situ hybridization (RNAscope, ACD)

Mice were anesthetized with isoflurane and rapidly decapitated. Brains were extracted and flash-frozen in isopentane on dry ice. Brains were stored at -80°C until sectioning. Prior to sectioning, brains were moved to -20°C and acclimated for 2–24 hours. Using a Leica CM8060 cryostat, 16µm slices were thaw mounted onto slides (Superfrost, Fisher) and stored at -80°C until assayed. RNAscope manual fluorescent multiplex v1, multiplex v2, and HiPlex assays were run as detailed in ACD technical manuals and previously reported (see https://acdbio.com/main-category/user-manuals)(4). For multiplex v1 and v2 experiments, markers were run separately (Figure 1b-d). For all in situ hybridization experiments, N represent animals with an approximately equal number of males and females. For v2 assays, we used Opal fluorophore reagent packs: 488, 520, 570, 620 and 780 (Akoya Biosciences). The following probes were purchased from ACD: Mm-Drd2-T1, Mm-Drd1-T2, Mm-Crh-T3, Mm-Otof-T4, Mm-Pcdh8-T5, Mm-Cacng5-T6, Mm-Tac2-T7, Mm-Crh-C1, CRE-C2, EGFP-C3, Mm-Vip-C2, Mm-Cck-C2, Mm-Nos1-C2, Mm-SST-C2, Mm-Npy-C2, Mm-Meis2-C2, Mm-Drd1-C2, Mm-Drd1-C3, Mm-Htr3a-C2, Mm-Calb2-C2, Mm-Chat-C3, Mm-Pdyn-C3, Mm-Drd2-C3, Mm-Penk-C3, Mm-Th-C2.

### Imaging and analysis

Tissue was imaged using a Keyence BZ-X710 fluorescence microscope or a Leica Stellaris 8 confocal microscope. For viral tracing experiments: 20x tile images of sagittal sections were taken, with additional 63x oil immersion, zoom 2 z-stacked and tiled images of regions of interest. Puncta were quantified using FIJI/ImageJ particle counter. For RNAscope experiments: 20x tiled images of the NAc were captured. For each run of RNAscope, capture settings were identical across animals. Settings were adjusted as needed between runs. DAPI expression was used to align images from separate HiPlex rounds as needed; this was done in Adobe Photoshop. For analysis, HALO image analysis platform (Indica Labs) was used. The NAc was either delineated by an atlas overlay using the ABBA FIJI/ImageJ plugin (BIOP, https://biop.github.io/ijp-imagetoatlas) or, for HiPlex experiments, outlined by hand. DAPI expression was used to delineate cells, and RNA puncta were quantified using the HALO FISH-IF module. For each probe, the threshold for minimum intensity, puncta size and number of puncta was titrated based on signal quality and expression pattern. The same analysis settings were applied across all sections from a single RNAscope run. For morphological analysis, FIJI/ImageJ plugins Dendritic Spine Counter and Neuroanatomy were used.

### Stereotaxic surgeries and viral vectors

Adult mice were quickly and deeply anesthetized using isoflurane (4% induction, 2% maintenance), and the skull leveled on a stereotaxic frame (Kopf, Model 1900). Viral particles were delivered using handmade 30g metal injectors connected to a 2uL syringe (Hamilton) with polyethylene tubing and delivered via pump (WPI microinjection syringe pump) at 100nL/min. For some experiments we also use glass sharp electrodes and Drummond Nanoject II microinfusion system. The injector was lowered to the appropriate depth and left to settle for one minute before beginning injection; following injection, the injector was left at depth for at least 5 minutes before the injector was slowly removed. For viral tracing experiments, either AAV5-hSyn-FLEx-GFP-2A-Synaptophysin-mRuby (Addgene plasmid #71760 gifted by Liqun Luo(5), virus packaged by UMN Viral Vector and Cloning Core) viral particles were delivered into the NAc (AP = 1.15, ML = ±1.1, DV = 4.7; mm relative to bregma; 250nL, 100 nL/min) of *Crh*^IRES-Cre/-^ mice. For optogenetic experiments, AAV5-EF1a-double floxed-hChR2(H134R)-EYFP-WPRE-HGHpA viral particles were delivered to the NAc (AP = 1.15, ML = ±1.1, DV = 4.7; mm relative to bregma; 250 nL, 100 nL/min). For some of the electrophysiology experiments, *Crh*^IRES- Cre/-^ mice were injected with AAV5-Ef1a-DIO-eYFP (AP = 1.2, ML = ±1.1, DV = 4.75; 300nL). For CRF conditional knockdown experiments, intracranial injections were done as described with either AAV8-CMV-eGFP-Cre-WPRE-SV40 (Addgene viral prep #105545-AAV8 gifted by James M. Wilson) or AAV8-CMV-eGFP-WPRE (Addgene viral prep #105530-AAV8 gifted by James M. Wilson), bilaterally into the NAc (AP = 1.2, ML = ±1.1, DV = 4.75; 300nL) of *Crh*^loxP/loxP^ mice. For fiber photometry experiments, intracranial injections were performed alongside fiber implantation. *Crh*^IRES-Cre/-^*; Ai14* mice were injected with AAV5-Syn-FLEx-GCaMP6s-WPRE (Addgene viral prep #100845-AAV5 gifted by Douglas Kim and GENIE Project(6)) unilaterally into the NAc (AP = 1.3, ML = ±1.3, DV = 4.7; 300nL), and a fiber-optic cannula (Doric, MFC_400/430-0.66_5mm_MF2.5_FLT) was placed dorsal to the injection bolus (DV = - 4.3). For electrophysiological experiments that assessed the effect of CRF knock-down on cellular function, we co-expressed AAV-hSyn-DIO-eGFP with AAV5-FLEX-SaCas9-U6-sg*Crh* or AAV5-FLEX-SaCas9-U6-sg*ROSA26* at a 1:4 ratio into the NAc (AP = 1.2, ML = ±1.1, DV = 4.75; 300nL, 100 nL/min). For co-expression of GCaMP6s with CRF CRISPR, *Crh*^IRES-Cre/-^ mice were injected with a mixture of AAV5-Syn-FLEx-GCaMP6s-WPRE and AAV5-FLEX-SaCas9-U6-sg*Crh* or AAV5-FLEX-SaCas9-U6-sg*ROSA26* at a 1:4 ratio into the NAc (AP = 1.3, ML = ±1.3, DV = 4.7; 300nL). Virus was injected bilaterally, but fiber placement was unilateral. Analgesic (carprofen, 5mg/kg s.c.) was given prior to and for 3 days after surgery.

### CRISPR viral vector generation

The single guide RNA (sgRNA) targeting *Crh* was generated following established protocols (Hunker et al., 2020)(7). The sgRNA was designed to target 153 base pairs downstream of the start codon for the prepropeptide. A sequence containing the BsaI overhang and sgRNA (Fwd: 5’ caccgCCTCAGCCGGTTCTGATCCGC 3’; Rev: 5’ aaacgCGGATCAGAACCGGCTGAGGC 3’) was cloned into the BsaI site of pAAV-FLEX-SaCas9-U6-sgRNA (Addgene #124844).

### Immunohistochemistry

Mice were intracardially perfused with PBS containing 10U/mL Heparin until the liver cleared (approximately 5 minutes) followed by 40 ml fixative solution (4% paraformaldehyde; 4% sucrose in 0.1M PB) at 3 ml/min. Brains were postfixed in fixative solution for 24 hrs, then incubated in 30% sucrose (in 0.1 M PB) for at least 48 h at 4 °C and sliced with a cryostat or microtome. 40µm floating sections were washed in PBS and then blocked for 1hr in 5% normal goat serum or donkey serum, 0.3% Triton-X in PBS at RT. Sections were then incubated in primary antibody for 12-18hrs at RT (Rat anti-Substance P, 1:400, EMD Millipore, Cat # Mab356; Chicken anti-TH, 1:1000, Aves Laboratories, Cat # TYH). Slices were incubated in goat or donkey Alexa Fluor secondary antibodies (1:500, Invitrogen) for 2 hr at RT. For TH immunoreactivity, slices were incubated in Goat anti-Chicken Biotin (1:500) for 2 hr RT and then Streptavidin-700 (1:500) for 1.5 hr RT. Slices were washed 3 x 10 min in PBS, then 2 x 10 min, mounted and coverslipped. Images (1024×1024) were acquired using a confocal microscope (Stellaris 8 Confocal microscope). Three to five confocal images were acquired per region (ventral pallidum, ventral midbrain) per mouse, with 5 mice total. Analysis was performed using ImageJ software on acquired images.

### Ex vivo electrophysiology

Adult (8-20 week-old) *Crh*^IRES-Cre/-^*; Ai14* mice were anesthetized using isoflurane, rapidly decapitated and brains quickly extracted for sectioning. Coronal 240 µm slices were cut on a vibratome (Leica VT1200S) in ice cold cutting solution. Slices were allowed to recover in warm (33°C) oxygenated artificial cerebrospinal fluid (ACSF) for 30-60 min before the recovery chamber was removed from the warm water bath and maintained room temperature for the remainder of the recording day. For electrophysiology experiments, slices were transferred to the recording chamber maintained at physiological temperature (30-32°C) in the recording chamber. A gigohm seal was achieved before achieving whole-cell configuration using electrodes (2-5MΩ) filled with a KMeSO_4_ internal solution. In whole-cell current clamp, passive membrane properties were recorded. A current-step protocol (-120 to +360pA, 40pA steps) was applied to examine the firing properties of these cells. For optogenetic experiments, sections from *Crh*^IRES-Cre/-^ mice injected into NAc with a cre-dependent channelrhodopsin (ChR2) expressing virus were prepared for either midbrain or ventral pallidum recordings. Midbrain sections were prepared as described above, cut horizontal from the ventral surface, and incubated in aCSF. For ventral pallidum sections, mice were anesthetized with pentobarbital (≥100mg/kg, i.p.) and transcardially perfused (30mL, 10mL/m) with warm aCSF (30-32°^C^) with kynurenic acid (3mM) before being rapidly decapitated and the brains extracted. Tissue was sectioned on a vibratome in warm aCSF with kynurenic acid, then incubated in aCSF with kynurenic acid, in a 32°^C^ water bath for 30 m before being moved to room temperature. For both midbrain and ventral pallidum recordings, sections were maintained in aCSF with NBQX and R-CPP (5µM each) for the duration of recording. Pipettes were filled with 50/50 CsMeSO_4_/CsCl internal solution with QX-314 (5mM) and 1% biocytin. Recordings were performed in whole-cell voltage clamp configuration with cells held at -80mV. Optogenetic stimulation was produced by a 470nm light emitting diode (LED; M470F3, ThorLabs) connected to a T-cube LED driver (LEDD1B, Thorlabs). The LED output was passed through furcation tubing (FT020, ThorLabs) which was stripped on one end to allow light to escape and be aimed at the recording site. The LED driver was connected to the patch clamp amplifier via BNC and light output controlled by TTL pulses. Paired pulse recordings were performed with two 5ms pulses (100ms inter-pulse interval) per sweep (15 sweeps, 10 s inter-sweep interval). In a subset of cells, Gabazine (5µM) was added to the bath following initial recordings and after 5 m of Gabazine treatment the paired pulse stimulation was run again. When assessing the effects of CRF deletion on cellular function, slices taken from *Crh^IRES-Cre/-^* mice transduced with cre-dependent eGFP/sgCrh or eGFP/sgRosa26 (control) were incubated in Gabazine and CGP55835 (5µM each) and cells recorded using a KMeSO_4_ internal solution. We first recorded spontaneous excitatory post-synaptic currents (sEPSCs) for 3 minutes in whole-cell voltage clamp, then switched to whole-cell current clamp to perform a current-step protocol. In adjacent slices, we used a 50/50 CsMeSO_4_/CsCl based internal in order to chloride load the cells. During the experiment, we incubated the slices in NBQX and R-CPP to isolate GABAergic spontaneous inhibitory post-synaptic currents (sIPSCs). We recorded cells for 5 minutes to allow for dialysis of the cell and analyzed the last three minutes. Data were acquired at 5 kHz and filtered at 1 kHz using Multiclamp 700B (Molecular Devices) or Double IPA (Sutter Instruments). Data were analyzed using pClamp (Molecular Devices) or SutterPatch (Sutter Instruments). In a subset of cells, 1% biocytin was included in the internal solution. These cells were fixed in 4% paraformaldehyde (PFA) for at least 24 hrs following recording, then post-hoc stained for streptavidin (Invitrogen, S32353) and imaged (Leica Stellaris 8, 63x). Measures of dendritic length, soma size, spine density, and dendritic intersections were made using FIJI/ImageJ. Solutions in mM. Cutting solution: 225 sucrose, 13.9 NaCl, 26.2 NaHCO_3_, 1 NaH_2_PO_4_, 1.25 glucose, 2.5 KCl, 0.1 CaCl_2_, 4.9 MgCl_2_, 4.9 MgCl_2_, and 3 kynurenic acid. ACSF: 124 NaCl, 2.5 KCl, 2.5 CaCl_2_, 1.3 MgCl_2_, 26.2 NaHCO_3_, 1 NaH_2_PO_4_ and 20 glucose (310-320 mOsm). KMeSO_4_ internal: 120 KMeSO_4_, 20 KCl, 10 HEPES, 0.2 K-EGTA, 2 MgCl_2_, 4 Na-ATP, and 0.4 Na-GTP (pH = 7.25, 290 mOsm). CsMeSO_4_/CsCl internal: 60 CsMeSO_4_, 60 CsCl, 10 HEPES, 0.2 EGTA, 8 NaCl, 2MgCl_2_, 2 Mg-ATP, 0.3 Na-GTP, and 10 phosphocreatine (pH = 7.24, ∼300 mOsm).

### Principal component analysis and hierarchical clustering

Analysis was performed on three minutes of stable, continuous recording following a five-minute equilibration period. For each cell, eight active and passive properties were extracted: membrane resistance, membrane capacitance, resting membrane potential, action potential duration, rheobase, subthreshold I-V slope, F-I curves and sag ratio. Passive properties (membrane resistance, membrane capacitance, and resting membrane potential) were measured immediately after establishing whole-cell access, while the remaining active properties were assessed after equilibration. Principal component analysis was performed on all eight electrophysiology parameters. Retaining principal components (PCs) with eigenvalues greater than 1, two PCs were selected that explained 78.71% of the variance. An unsupervised clustering approach was used to examine whether neurons grouped naturally into the three clusters observed by experimenters. Agglomerative hierarchical clustering was implemented in R (version 4.3.1) on all eight electrophysiology parameters; Ward’s linkage method provided the strongest clustering structure. Experimenter-determined cell type designations were included as labels on the resulting dendrogram to allow for assessing the degree of concordance between experimenter-determined cell types and hierarchical clustering results.

### Operant training

Mice were allowed to recover from surgery for 21-28 days prior to behavioral testing. One week prior to training, mice were gradually food restricted to 85-90% body weight. Mice were placed in modular chambers (Med Associates) with two illuminated nose poke ports and a trough pellet receptacle. Mice were first trained on a fixed ratio 1 (FR1) schedule with one active and one inactive nose poke port. Active and inactive sides were counterbalanced across groups. Sessions lasted 30 min or until 50 pellets were earned, whichever occurred sooner. A rewarded nose poke triggered a pellet to be dispensed into the trough receptacle and a pure tone to play for 1 s. For the first 3 days of training, the pellet receptacle was baited with between one and three chocolate pellets (Bio-Serv, F05301) and both nose poke ports were baited with dust from a crushed chocolate pellet. During CRISPR fiber photometry experiments, some mice had trouble adjusting to the patch cord. For mice failing to engage in the task (defined as fewer than 20 rewards earned) after 5 days, the nose poke ports were re-baited for two days. Mice were run on an FR1 schedule until criteria, at which point they were moved to FR3. For *Crh*^loxP/loxP^ cKO experiments, the criterion for phase completion was 3 consecutive days with at least 30 earned rewards and 75% discrimination of the active lever. For fiber photometry experiments, criterion for phase completion was 4 consecutive days with 30 earned rewards and 75% discrimination, to allow for at least two recorded days per phase per mouse. For *Crh*^loxP/loxP^ cKO and naïve fiber photometry experiments: after reaching criteria at FR3, mice were tested in a 1-day progressive ratio task. The progressive ratio task was based on a protocol modified for mice by Johnson et al. (2022(8)), but with a maximum session time of 30 min. For CRISPR experiments: after reaching criteria at FR3, mice were run through a 5 day reversal test, modified from Whitehouse et al. (2017)(9). On reversal days, the active and inactive nose ports were switched (such that the previously active port became inactive and vice-versa) and mice were run on an FR1 schedule with the same parameters as FR1 training (30 min or 50 reward max). Criteria were not considered for reversal days: mice were run for 5 consecutive days regardless of performance. Three mice were excluded from fiber photometry experiments due to non-responding after 10 days. For *Crh*^flox/flox^ cKO studies, four mice (two control, two cKO) were excluded due to poor viral expression. In CRISPR fiber photometry experiments, three mice were dropped for non-responding (one control; two sgCrh) and one mouse was dropped after losing a head-cap (control).

### Fiber photometry

Mice were trained on an operant task as described above. During training, mice were attached to a 400µm core fiberoptic patchcord (Doric) by a black covered zirconia mating sleeve (Doric) to reduce light leakage. Mice were attached every day of training, but recordings were only performed on alternating days to prevent bleaching. A Doric/TDT fiber photometry system was used to record fluorescence. This system included: integrated fluorescence mini-cube (Doric), femtowatt photoreceiver module (Doric, 2151), LED driver (Doric, LEDD_2), two connectorized LEDs (Doric, CLED_405 and CLED_465), patch panel (Tucker Davis Technologies, PP24), fiber photometry processor (TDT, RZ5P) and high-performance computer workstation (TDT, WS4) with Synapse software (TDT) to control the LEDs and collect data. A 10 Hz lowpass filter was applied at the time of collection, and LED outputs were calibrated daily to 40µW output at the patchcord tip. TTLs generated by Med-PC (Med Associates) were used to time-lock recordings with behavioral events (nose poke, entry into the magazine, reward dispensed). Analysis of photometry signals was done using the Python toolbox GuPPy, created by the Lerner Lab(10). A zero-phase moving average linear digital filter was used with a 100 data point window. An isosbestic control channel (405 nm) was used to filter out artifacts, and a standard z-score was calculated. The Med-PC generated timestamps were used to align z-scores from behavioral events, and these z-scores were first averaged by session then by mouse.

Following fiber photometry recordings, mice were anesthetized with pentobarbital (Fatal-Plus; 100mg/kg, i.p.), and transcardially perfused with ice cold 1x PBS (8 mL) followed by 4% PFA (50 mL). Brains were post-fixed in 4% PFA for 24 hours, then sunk in 30% sucrose in 1x PBS for 48-72 hours. On a microtome (ThermoFisher Microm HM 450), tissue was sectioned in 40 μM coronal slices free floating in 1x PBS with 0.01% sodium azide. Sections containing NAc were mounted onto slides and fiber placement was confirmed by microscopy.

### Sucrose preference

Mice were individually housed for the duration of sucrose preference testing. Each cage was given two identical water bottles, which were weighed every 24 hours (same time of day) and placed back into the opposite side of the cage after weighing. On days 1-4, both water bottles were filled with water in order to habituate the animals. On days 5-7, one of the two bottles contained 1% sucrose solution. The sucrose bottle side was counterbalanced across groups and switched daily after weighing. A dummy cage containing water/water on days 1-4 and sucrose/water on days 5-7 was also weighed to control for non-drinking loss of liquid and ensure that water and sucrose had comparable ambient loss. Sucrose preference was calculated for each mouse each day by calculating the percentage of daily intake that was sucrose [sucrose/(sucrose+water)].

### Open field exploration

Open-field locomotion was measured in large custom made acrylic chambers (50 x 50x 40 cm) for 60 minutes using video monitoring and Noldus Ethovision software.

### Quantitative polymerase chain reaction

The nucleus accumbens was rapidly dissected from *Crh*^loxP/loxP^ mice transduced with AAV5-CMV-eGFP or AAV5-CMV-Cre-eGFP and stored in the -20 freezer in RNAlater. Primers used for PCR of the *Crh*^loxP^ (*Crh-*Forward: TTTATGGCCTTCCTCGTTAG; *Crh*-Reverse: TCTGTCGTTACTATGGCCTG). Total RNA was purified using RNeasy Micro (Qiagen). RNA was quantified using a Nanodrop and cDNA was synthesized using an Invitrogen kit. qPCR for RNAs for *GAPDH* and *Crh* were run using TaqMan Gene Expression Assays (Thermofisher, Cat#4331182). We determined relative mRNA expression of the endogenous control gene GAPDH and *Crh*, respectively. Quantitative PCR (qPCR) was performed using TaqMan Fast Polymerase (Applied Biosystems) in a StepOnePlus Real-Time PCR system. Cycling conditions were as follows: initial hold at 95°C for 20 s; 40 cycles of step 1 (95°C for 1 s); and step 2 (60°C for 20 s). Samples were run in quadruplicate, and negative controls were run in parallel. cDNA synthesis and qPCR experiments were repeated three times. Relative quantification was calculated using the ΔΔCt method (StepOne System Software, Applied Biosystems).

### Corticosterone ELISA

Mice were killed by rapid decapitation and trunk blood was collected. Trunk blood was allowed to coagulate for 30 min at room temperature. Samples were then centrifuged for 10 min at 2000 x g. Serum was collected and stored at -20°C. ELISA was run according to manufacturer instructions (Enzo, ADI-900-097). In brief, samples were pipetted into the assay plate, then buffer, blue and yellow conjugates were added. The plate was incubated for 2 hrs with agitation (up to 500 rpm). Wells were then emptied and washed three times before pNpp substrate was added and incubated for 1 hr. Finally, stop solution was added and optical density measured on a microplate reader (SpectraMax, Molecular Devices). Corticosterone standards (20000, 4000, 800, 160, 32 pg/mL) were run alongside and used to fit a 4 parameter logistic curve.

## Supplemental Discussion

### Is the NAc more like the dorsal striatum or the extended amygdala?

In the bed nucleus of the stria terminalis (BNST), CRF neurons can be segregated into three groups based on their electrophysiological properties. Moreover, the electrophysiological phenotypes we identify in the NAc are similar to what has been reported for CRF+ neurons in the BNST. It remains a point of debate whether the NAc is part of the striatum, part of the extended amygdala or its own unique region.

Considering that at least portions of the NAc (NAc shell) are considered by some to be part of the extended amygdala(11), it is worth considering the electrophysiological properties of BNST CRF neurons. In 2013, Winder and colleagues identified two types of BNST CRF neurons that had low-threshold spiking properties and one type that displayed canonical SPN-like properties(12), similar to our results here. It is possible that developmentally, NAc CRF neurons retain the electrophysiological features of their BNST counterparts.

### Alternative interpretation of CRF regulation of reward learning: “Throw caution to the wind…”

An alternative explanation to our behavioral data is that CRF, as a stress-associated neuropeptide, is suppressing food acquisition and consumption in the context of an instrumental learning task during a period in the task where the environment is most novel and most stressful. The antagonistic relationship between stress and feeding (i.e., stress-induced hypophagia) in rodents has been well described(13). Indeed, a classic stress-induced anxiety measure is novelty-induced suppression of feeding. Stress-induced hypophagia is often attributed in part to CRF neurons in the hypothalamus modulating feeding-associated neurons in the arcuate nucleus(13). However, it is possible that in the context of reward learning, CRF from this sparse population of NAc neurons constrains initial acquisition in the face of high novelty by tempering the “go” signal from direct pathway SPNs, which is thought to potentiate and reinforce behaviors(14, 15). CRF is generally an excitatory transmitter; both CRF receptors couple to either Gαs or Gαq and, therefore, would potentially counteract inhibitory GABA release from SPNs(16). This may be the case particularly when both the effort and reward payoff is low. Bryce and Floresco demonstrated that intra-NAc microinfusion of CRF reduces action-outcome performance under low effort, low reward conditions (like our task), but increases the performance under high effort, high reward conditions(17). Unfortunately, there is a dearth of information on the function of CRF in the ventral pallidum and ventral midbrain, particularly in the context of reward processing, so this interpretation is fairly speculative.

## Supplemental Figures

**Figure S1, related to Methods.**
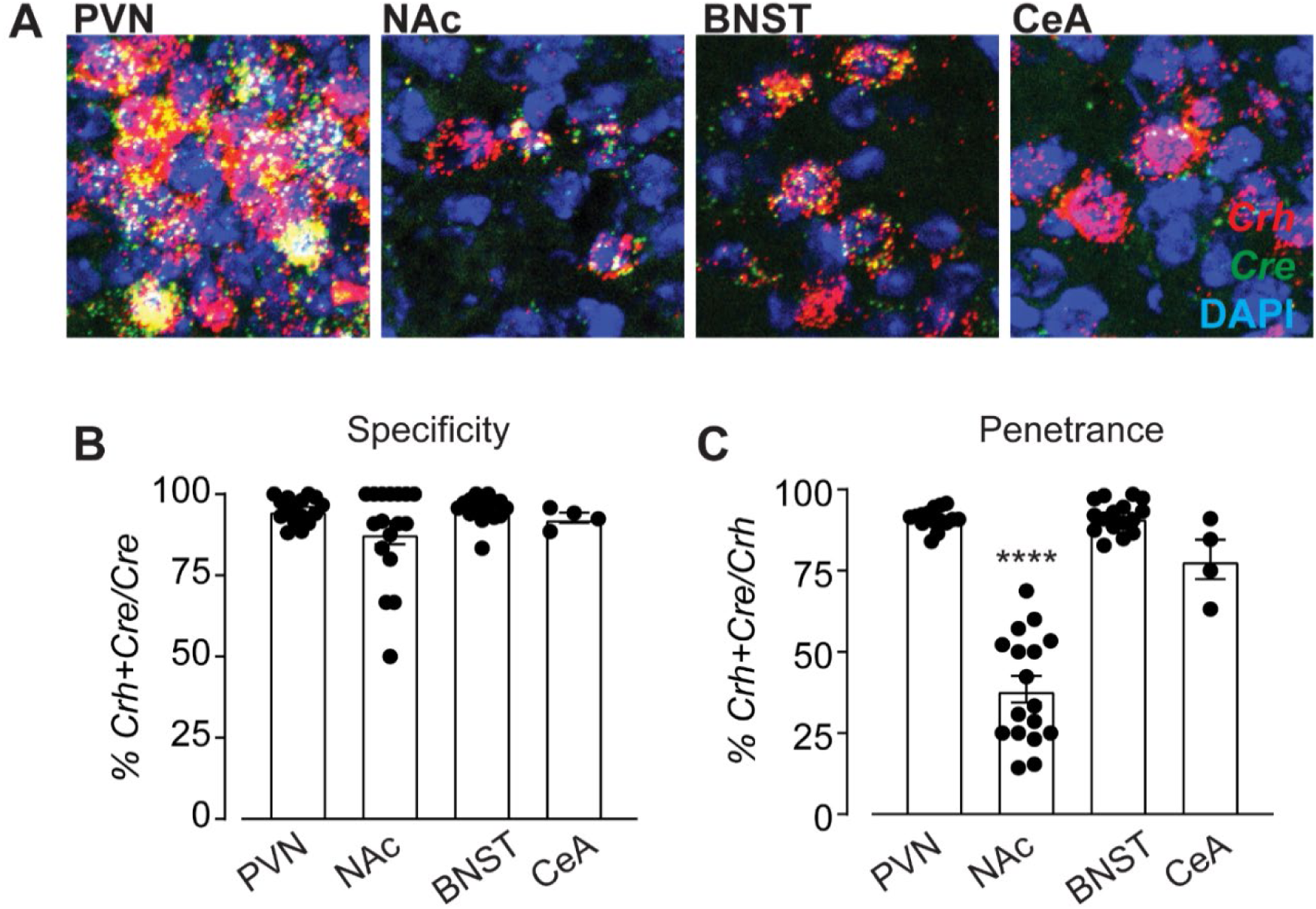
Validation of CRH^IRES-Cre/-^ mouse line. (A) Representative images of in situ hybridization expression from paraventricular nucleus of the hypothalamus (PVN), nucleus accumbens (NAc), bed nucleus of the stria terminalis (BNST) and central amygdala (CeA) from *Crh*^IRES-Cre/-^ animals for *Crh* (red) and *Cre* (green) mRNA with DAPI counterstaining. (B) Specificity of the *Crh*^IRES-Cre/-^ mouse line was assessed across regions as the percentage of *Cre*+ cells that were also positive for *Crh* mRNA. There were no significant differences across regions, with high specificity in all regions (PVN: 95 ± 1%; NAc: 88 ± 4%; BNST 93 ± 2%; CeA 95 ± 4%; one-way ANOVA: F_3,49_ = 2.276, p = 0.09, N = 4-16). (C) Penetrance of the *Crh*^IRES-Cre/-^ mouse was assessed across regions as the percentage of *Crh*+ cells that were also positive for *Cre* mRNA. Penetrance was high in the PVN, BNST and CeA, but was significantly reduced in the NAc (PVN: 91 ± 1%; NAc: 38 ± 4%; BNST 92 ± 1%; CeA 78 ± 6%; one-way ANOVA with = F_3,49_ = 93.75, p < 0.0001, Tukey’s post-hoc t-tests of NAc vs. PVN,BNST and CeA, N = 4-17). (Data are represented as mean ± SEM.) (****p < 0.0001).

**Figure S2, related to Figure 1.**
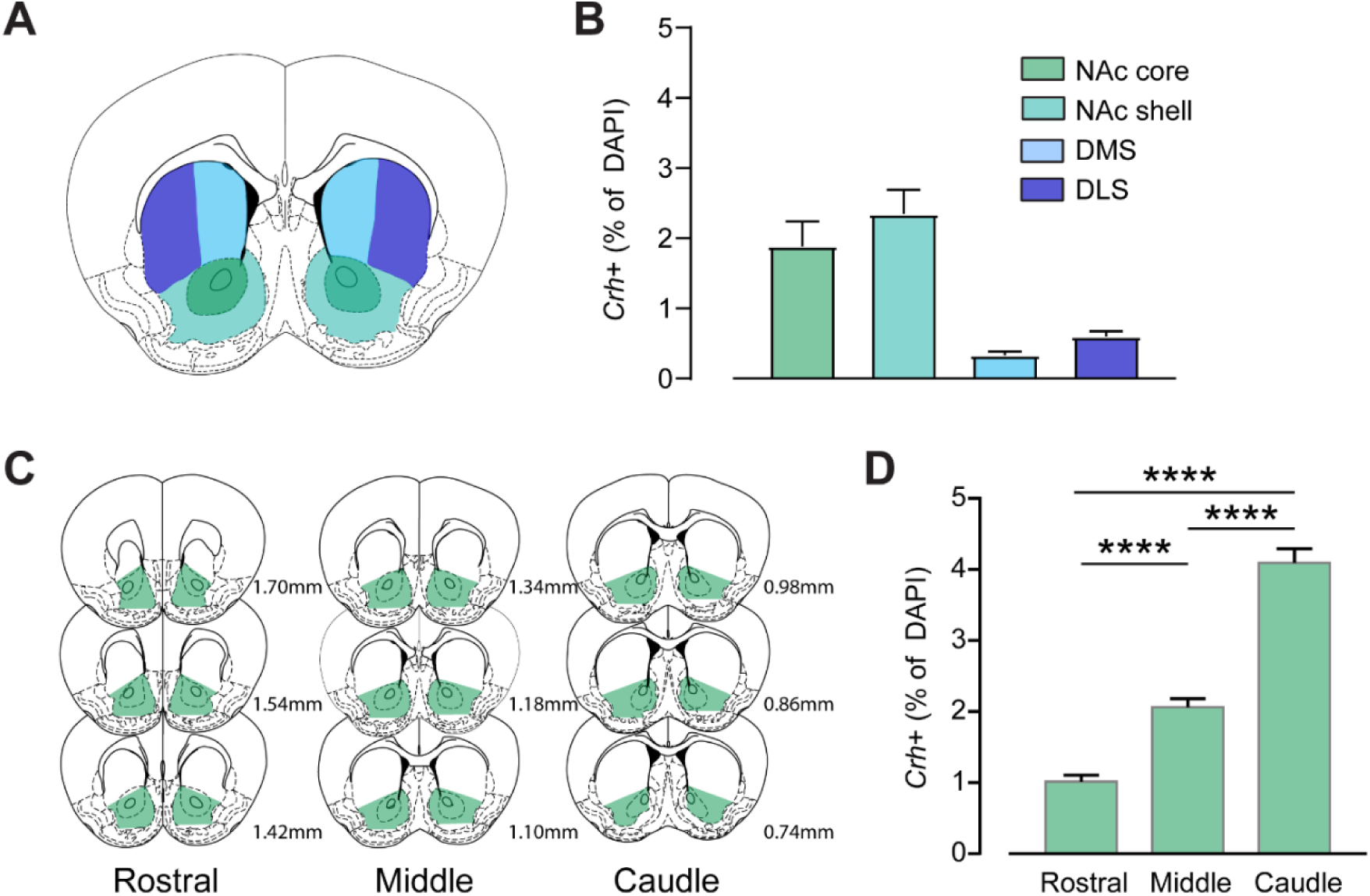
Crh mRNA expression pattern in striatum. (A) Schematic of compartmentalization of dorsal medial striatum (DMS, light blue), dorsal lateral striatum (DLS, dark blue), nucleus accumbens core (NAc core, dark green), and nucleus accumbens shell (NAc shell, light green). (B) Quantification of in situ hybridization for *Crh* mRNA as percent of total DAPI+ cells in striatal compartments. *Crh+* cells were more abundant in the NAc (NAc core: 2.42 ± 0.25%; NAc shell: 2.58 ± 0.27%) than in the dorsal striatum (DMS: 0.47 ± 0.05%; DLS: 0.61 ± 0.06%) (DS vs. NAc, unpaired t-test, p = 0.0135; N = 6 mice). (C) Schematic of rostral, medial and caudal delineations relative to bregma for NAc sections. (D) *Crh* mRNA expression as percent of total DAPI+ cells in NAc across the rostral-caudal axis. NAc *Crh*+ expression followed a rostral to caudal gradient pattern. Rostral NAc contained the fewest Crh+ cells. Middle NAc contained significantly more than rostral NAc. Crh was the most abundant in caudal NAc compared to both rostral and middle compartments (rostral: 1.013 ± 0.091%; middle: 2.061 ± 0.121%; caudal: 4.092 ± 0.198%; one-way ANOVA: F_2,140_ = 99.40, p < 0.0001, post-hoc Tukey’s test comparing groups, N =29-62 sections collected from 12 mice). (Data are represented as mean ± SEM.) (****p < 0.0001).

**Figure S3, related to Figure 2.**
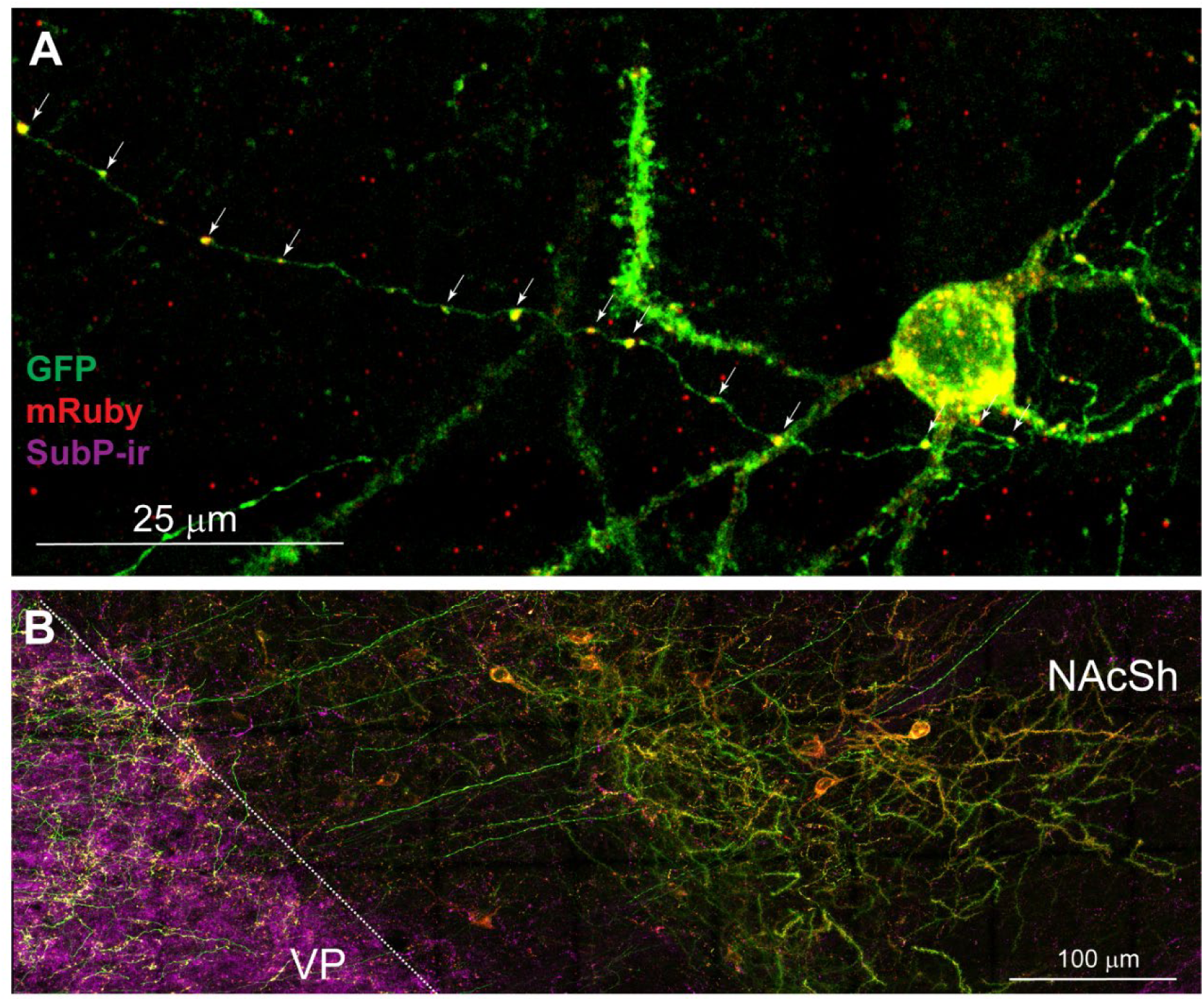
Additional images of NAc^CRF^ projections within the NAc and to the VP. High power confocal images from a *Crh*^IRES-Cre/-^ mouse with AAV5-hSyn-FLEx-GFP-2A-Synaptophysin-mRuby injected into the NAc. (A) A transfected NAc^CRF^ putative SPN shows that GFP (green) can be seen in the cell body and along processes. mRuby (red) was found on varicosities, suggesting *en passant* release sites. (B) Substance P immunoreactivity (magenta) used to delineate VP. A cluster of GFP+ CRF cells can be seen at the boundary between NAc and VP.

**Figure S4, related to Figure 3.**
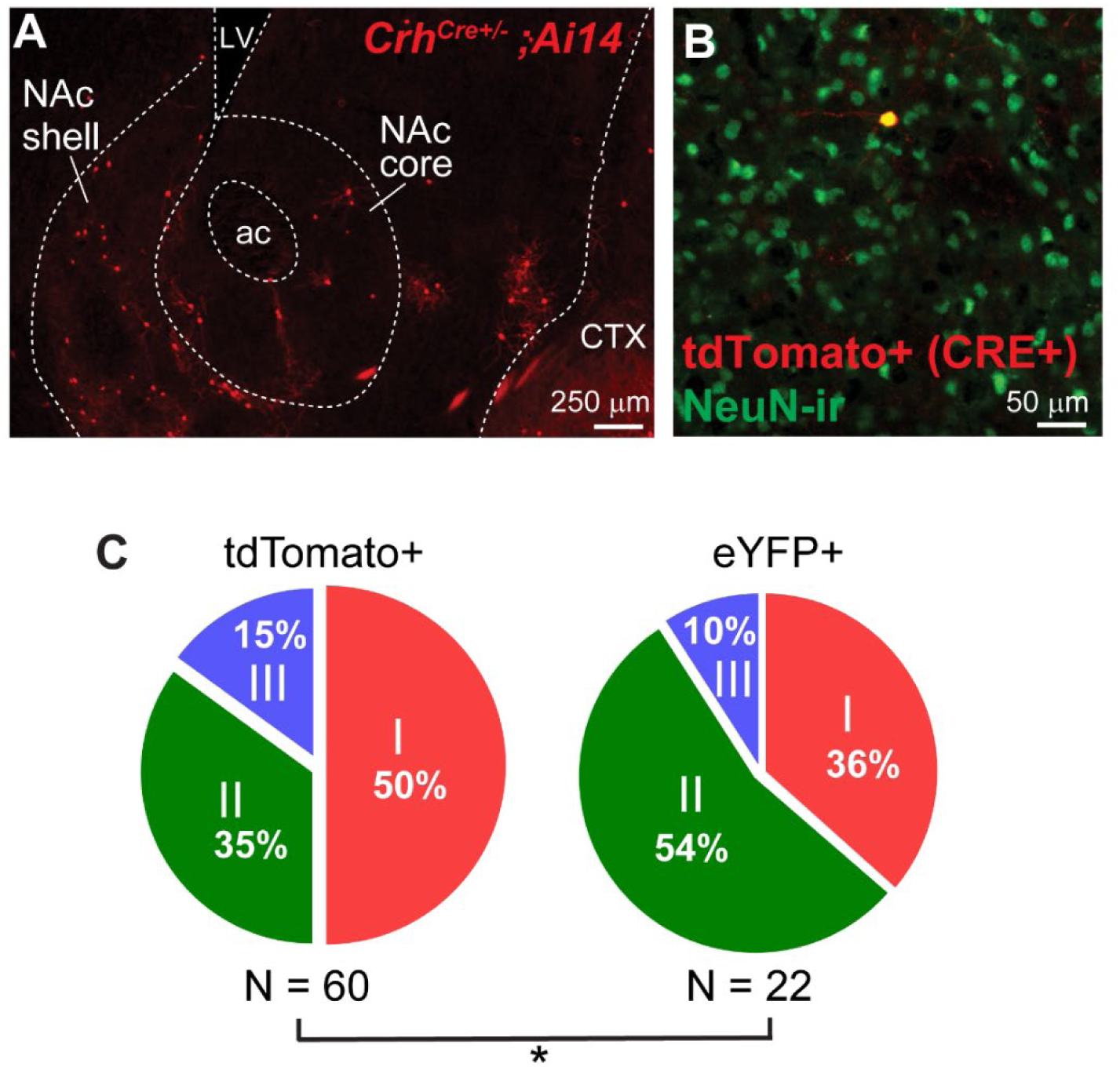
Proportions of NAc^CRF^ neuron electrophysiological subtypes recorded from Crh^IRES-Cre/-^ x Ai14 reporter cross versus viral transduction of AAV5-hSyn-DIO-eYFP into Crh^IRES-Cre/-^ NAc. (A) Representative image from a *Crh*^IRES-Cre/-^; Ai14 reporter mouse of Cre-dependent tdTomato (red) expression in the NAc. (B) High magnification image from a *Crh*^IRES-Cre/-^; Ai14 mouse with Cre-driven tdTomato (red) expression co-localized with a NeuN (green) positive cell body. (C) Relative proportions of subtypes from all NAc^CRF^ neurons recorded from either *Crh*^IRES-Cre/-^; Ai14 reporters (tdTomato+; left) or *Crh*^IRES-Cre/-^ mice injected with AAV-DIO-eYFP viral particles (eYFP+; right). There was a shift in the distribution of CRF subtypes depending on which method was used, suggesting that there may be a developmental shift in the NAc^CRF^ population (Chi-squared test, p = 0.0255).

**Figure S5, related to Figure 3.**
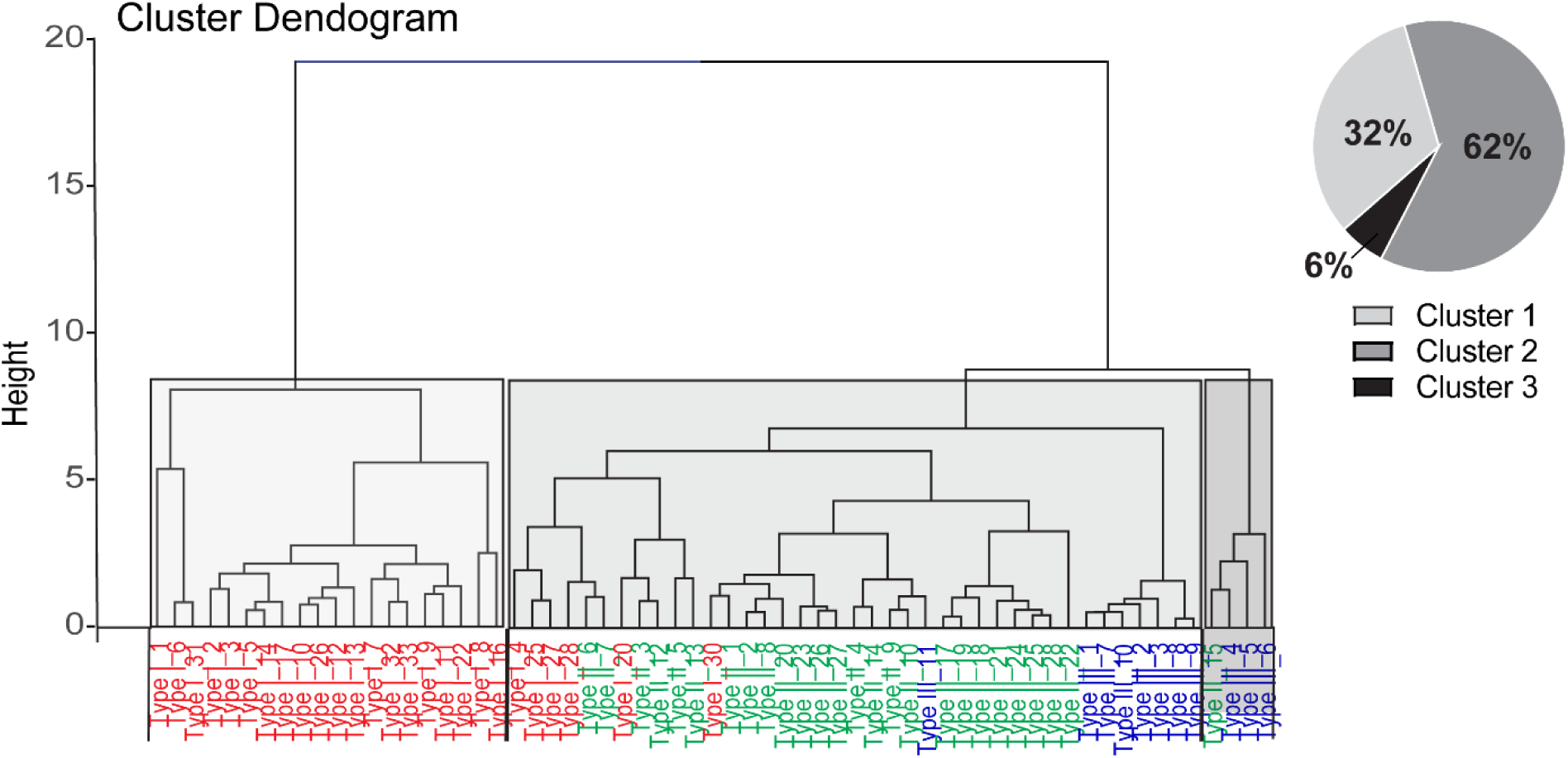
Hierarchical clustering of NAc^CRF^ cells based on electrophysiological properties. Hierarchical clustering of CRF cells based on active and passive electrophysiological components. While three clusters were sorted using this method, the experimenter defined subtypes (Type 1, red; Type 2, green; Type 3, blue) were not precisely replicated. The largest cluster, Cluster 2 (62%) is mostly composed of Type 2 neurons but also contains a number of Type 1 and Type 3 cells. On the other hand, Cluster 1 (32%) is made up entirely of Type 1 neurons, while Cluster 3 (6%) is composed of Type 3 cells with one exception.

**Figure S6, related to Figure 3.**
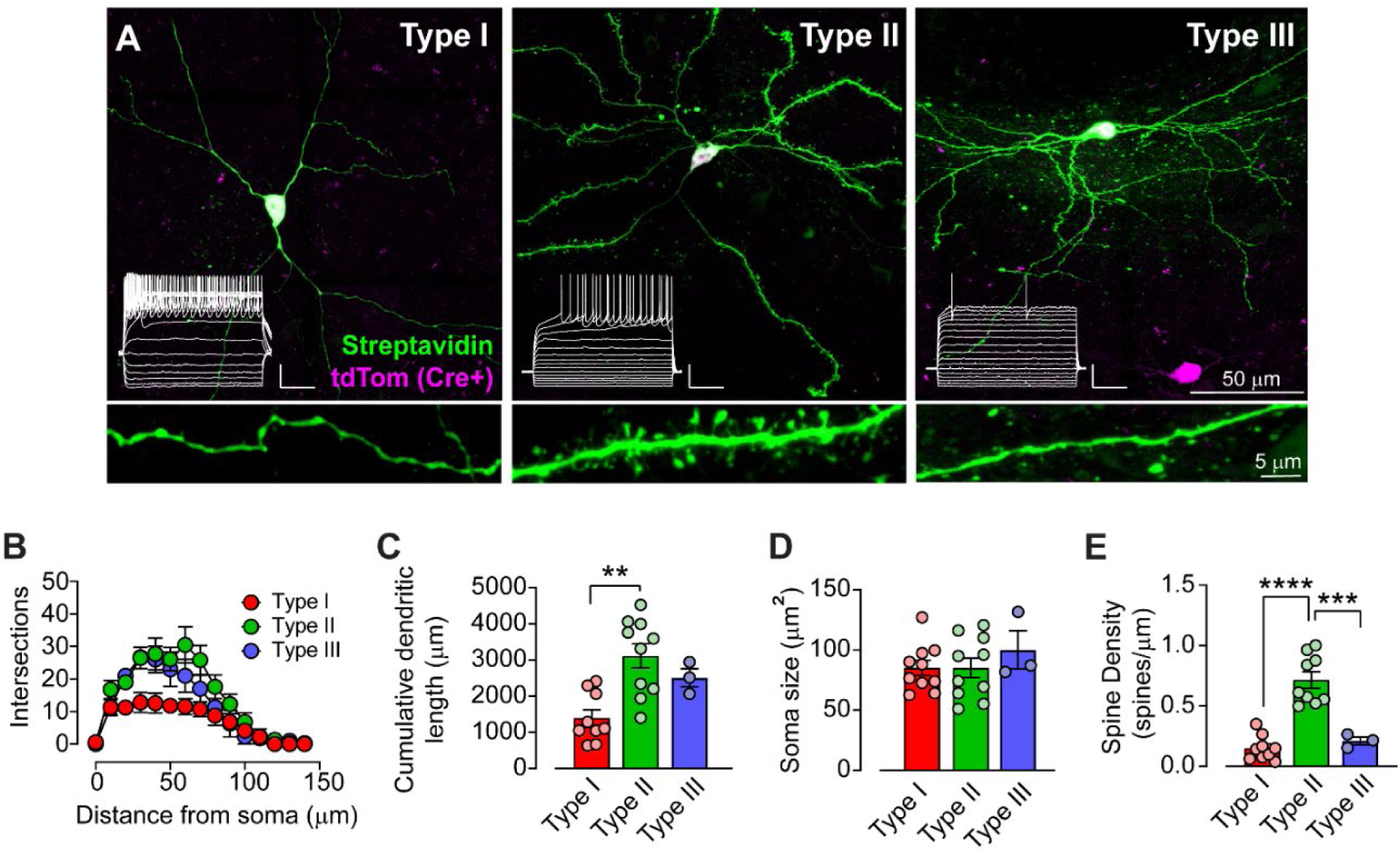
Accumbal CRF subtype confers discrete morphological profile. (A) Example images of NAc^CRF^ neuron electrophysiological subtypes recorded from *Crh*^IRES-Cre/-^; Ai14 mice. tdTomato (pink) expression driven by Cre recombination was used to identify CRF+ cells which were filled with biocytin during recording and post-hoc stained for streptavidin (green). (inset) Example traces from pictured cells in response to a step-current protocol. (B) Sholl analysis of dendritic intersections as a function of distance from the soma across NAc^CRF^ cell subtypes. Type I (red; N_cells_ = 9) cells had significantly fewer intersections compared to Type II (green; N_cells_ = 10) and Type III (blue; N_cells_ = 3) cells (Two-way RM ANOVA, distance x type interaction, F_28,266_ = 2.501, p < 0.0001). (C) Analysis of the total dendritic length across CRF cell subtypes. Type I had significantly smaller cumulative dendritic length compared to Type II or Type III cells. (One-way ANOVA, F_2,19_ = 9.413, p = 0.0014, Tukey’s post-hoc test, Type I vs 2, p = 0.0010). (D) Comparison of soma size between CRF cell subtypes, with no significant difference between groups. (One-way ANOVA, F_2,20_ = 0.5341, p = 0.5943). (E) Quantification of dendritic spine density across subtypes of CRF cells. Type II cells had significantly increased spine density compared to Type I and Type III cells. (One-way ANOVA, F_2,18_ = 33.97, p < 0.0001, Tukey’s post-hoc test, Type I vs II, p < 0.0001; Type II vs. Type III, p = 0.0003). (Data are represented as mean ± SEM; scale bar: 25mV, 100ms) (**p = 0.0010, ***p = 0.0003, **** p < 0.0001, one-way ANOVA.)

**Figure S7, related to Figure 3.**
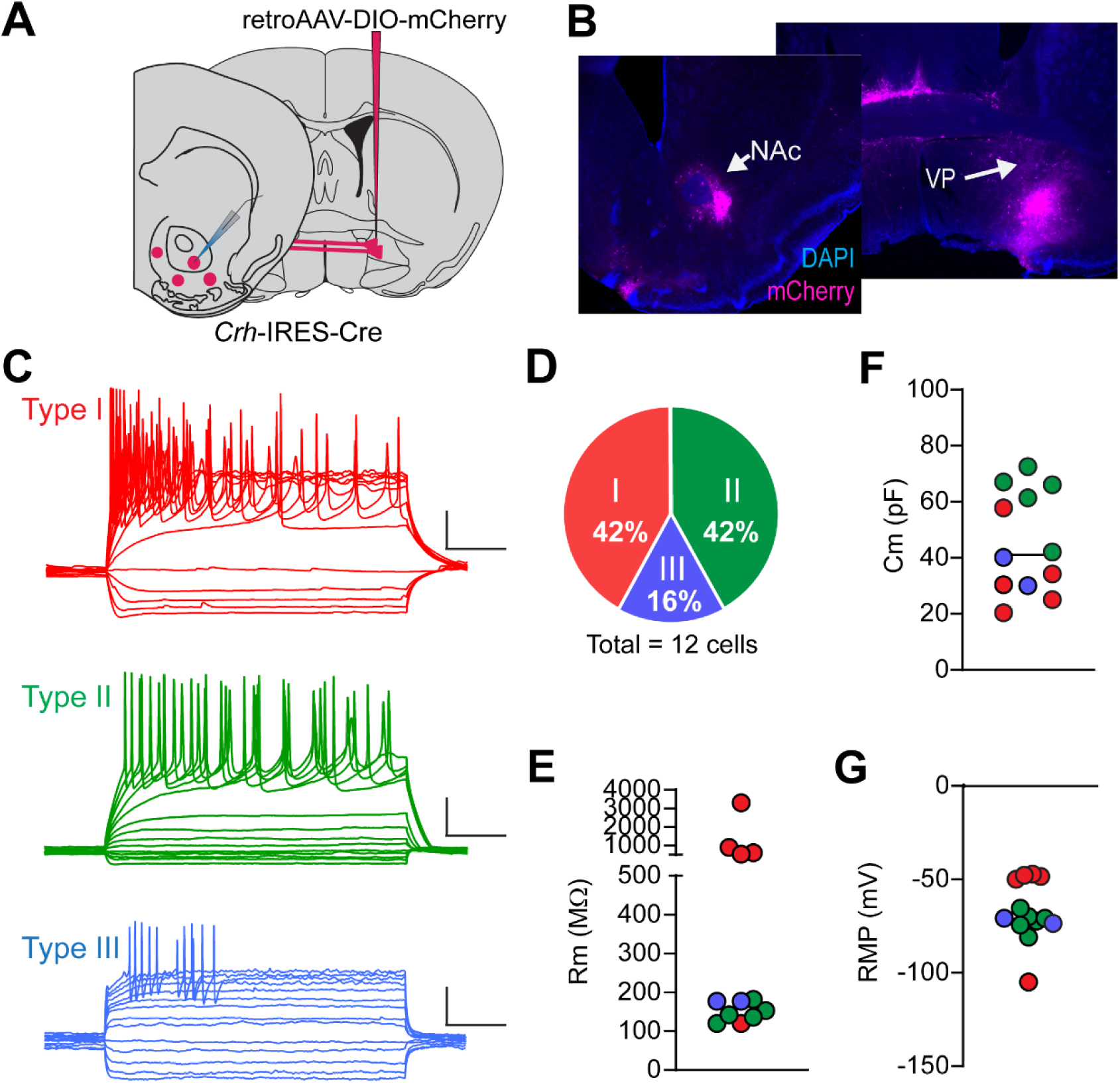
VP-Projecting NAc^CRF^ neurons are composed of all three CRF subtypes. (A) Schematic of the experimental preparation. *Crh*^IRES-Cre/-^ mice were injected with rgAAV-hSyn-DIO-mCherry or rgAAV-hSyn-DIO-eGFP viral particles into the VP and ex vivo recordings were performed from fluorescently labeled neurons in the NAc. (B) Representative images from the injection site (VP) and recording location (NAc) of mCherry expression (pink). (C) Example traces from current-step recordings of VP-projecting NAc^CRF^ neurons. Individual cells were sorted by subtype based on prior classifications. (D) Relative distribution of subtypes from VP-projecting NAc CRF cells recorded. All three subtypes were observed, suggesting that all subtype populations are at least partially projection cells. The relative proportion of each subtype was similar to observations from recordings of all CRF NAc neurons (42% Type I, 42% Type II, 16% Type III, N_Cells_ = 12). (E) Membrane resistance of NAc^CRF^ ◊ VP cells, with resistance of Type I cells (red) being higher on average, as previously observed. (F) Capacitance and (G) resting membrane potential of VP-projecting NAc CRF+ cells recorded.

**Figure S8, related to Figure 5.**
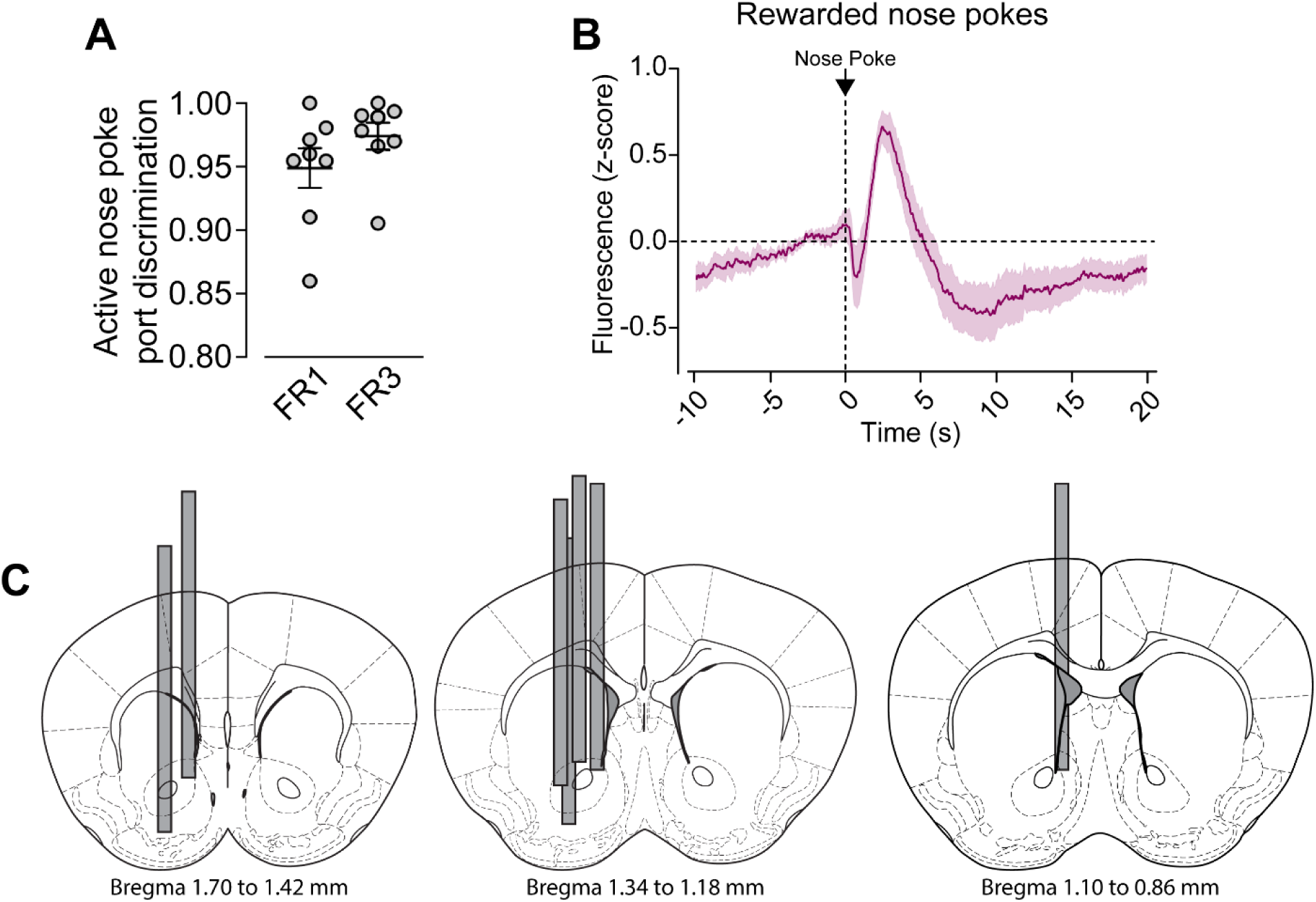
Additional behavior and recording analysis from fiber photometry recordings. (A) Discrimination of the active nose port [active pokes/(active+inactive pokes)] during FR1 and FR3 training, averaged by mouse. (B) Average fluorescence (z-score) aligned to TTLs elicited by a rewarded nose poke from FR1 and FR3 sessions. An immediate depression of activity was apparent following reward delivery (∼0 to 1s), followed by an increase in fluorescence beginning ∼1s after reward delivery, which likely corresponds to head entry into the magazine. (C) Representation of fiber placements from all mice included in fiber photometry analysis.

**Figure S9, related to Figure 6.**
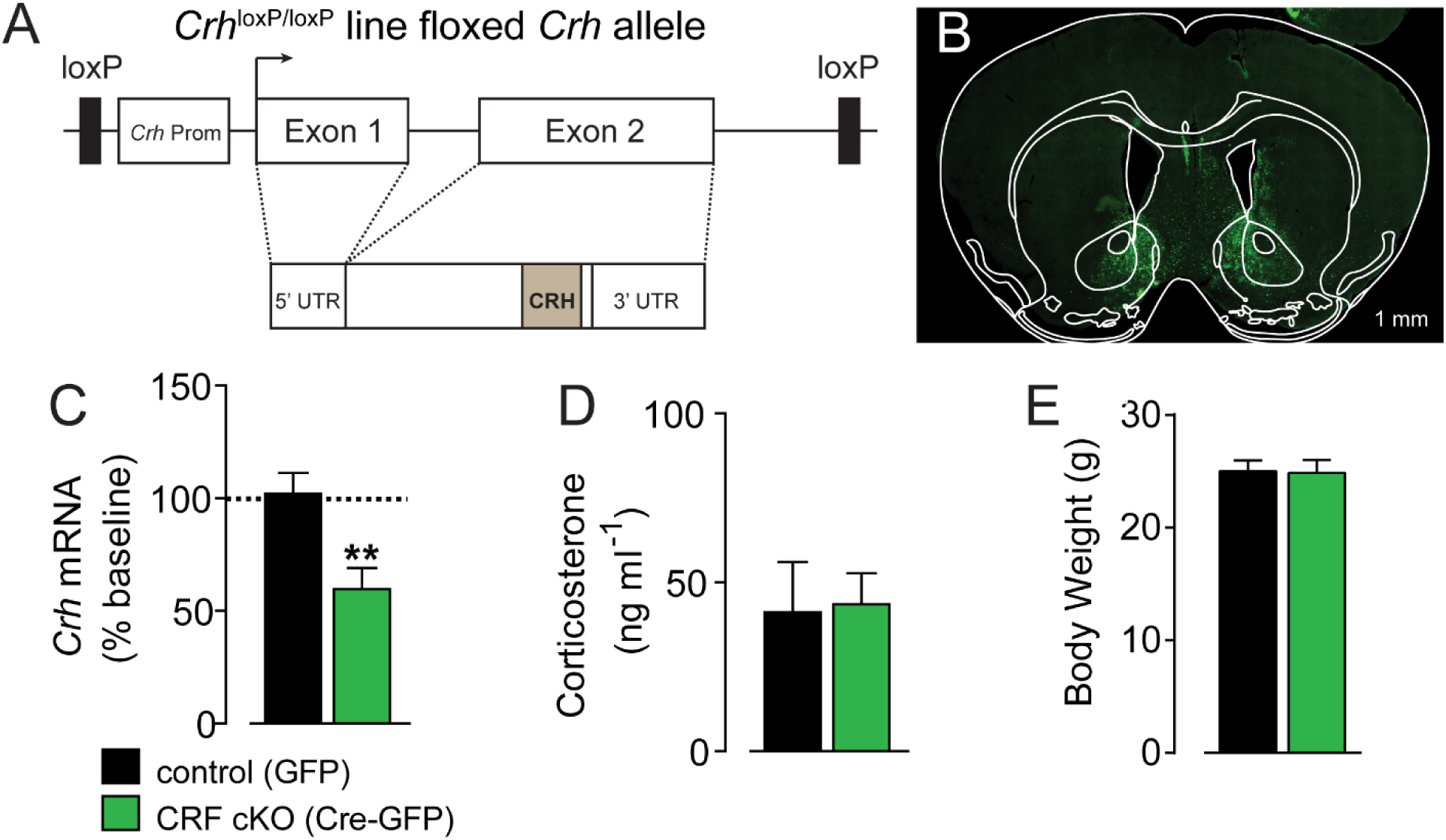
Validation of NAc CRF conditional knock-out. (A) Schematic of floxed *Crh* locus from the *Crh*^loxP/loxP^ mice used to generate conditional knock-outs. (B) Representative image from a *Crh*^loxP/loxP^ mouse injected bilaterally with AAV5-CMV-Cre-GFP (CRF cKO) into NAc. (C) *Crh* mRNA in NAc of control (AAV5-CMV-GFP; black) or CRF cKO (green) mice after viral injection. RT-qPCR was used to quantify *Crh* mRNA. All values were normalized to average control *Crh*. cKO mice had significantly reduced *Crh* (normalized control: 1.09 ± 09, N = 6; normalized CRF cKO: 0.59 ± 0.09, N = 7, unpaired t-test, p = 0.0084). (D) Corticosterone measured from trunk blood of control and CRF cKO mice. There was no impact of NAc CRF cKO on plasma corticosterone from trunk blood. (E) Body weight of control and CRF cKO mice. There was no difference between groups. (Data are represented as mean ± SEM.)

**Figure S10, related to Figure 6.**
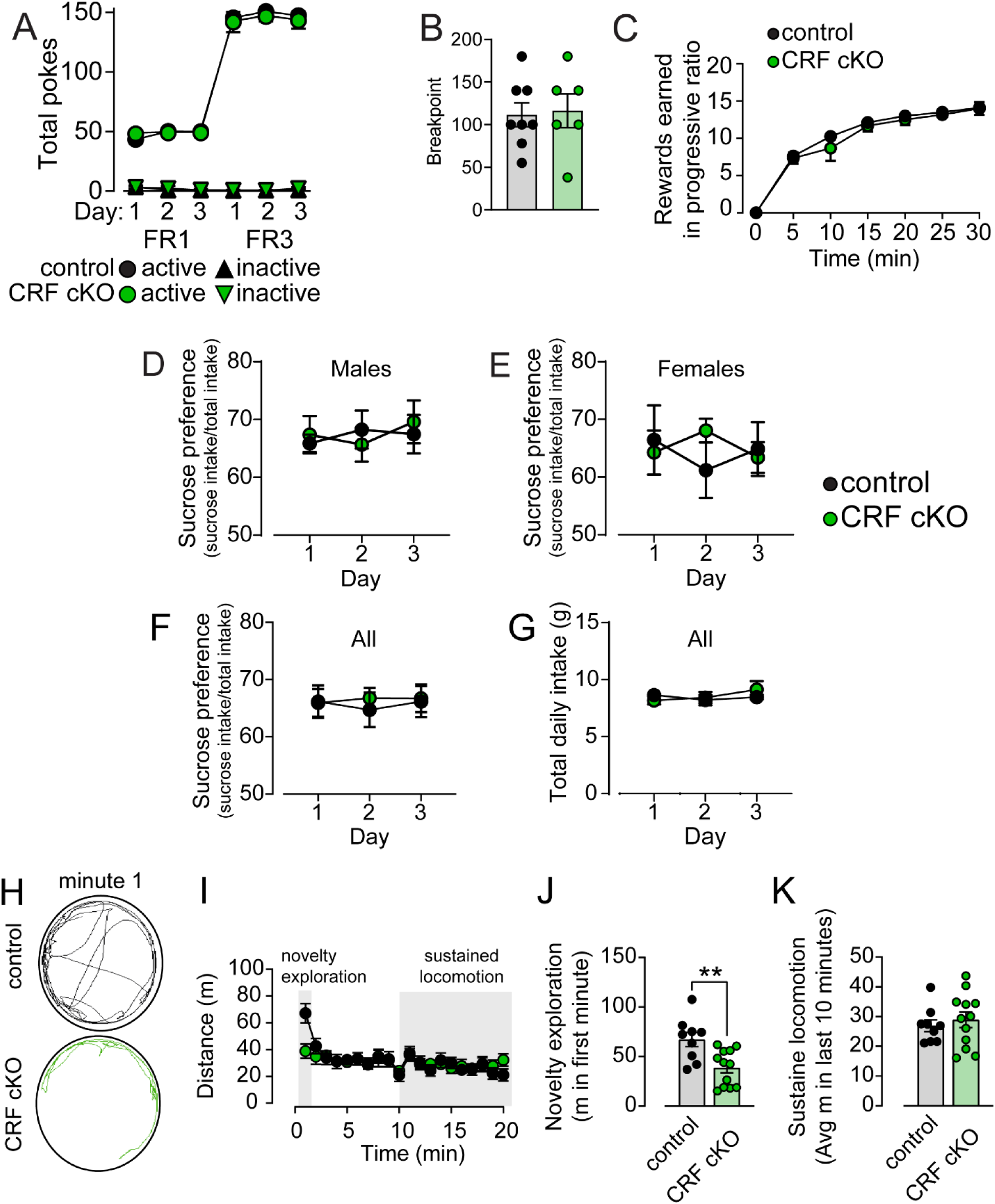
Additional behavior data from CRF cKO experiments. (A) Daily total pokes to active (circles) and inactive (triangles) nose ports from control (black) and CRF cKO (green) mice. There was no difference in total pokes between groups during fixed ratio 1 (FR1) or fixed ratio 3 (FR3) training (Mixed effects analysis. Active pokes: main effect of day: F_2.080,24.61_ = 768.6, p < 0.0001; group: F_1,12_ = 0.2208, p = 0.6469; day x group: F_6,71_ = 0.6918, p = 0.6569) (Inactive pokes: main effect of day: F_2.371,28.06_ = 3.936, p = 0.0254; group: F_1,12_ = 0.7424, p = 0.4058; day x group: F_6,71_ = 0.4281, p = 0.8579). (B) Breakpoint values for control and CRF cKO mice from the progressive ratio task. There was no difference in breakpoint between groups (unpaired t-test, p = 0.8451). (C) Number of rewards earned over time in the progressive ratio task. There was no difference between control and CRF cKO mice in rewards earned over time in the progressive ratio task (main effect of time: F_6,72_ = 225.2, p < 0.0001; group: F_1,12_ = 0.3674, p = 0.5557; group x time: F_6,72_ = 0.6322, p < 0.8562). (D-G) To assess whether conditional deletion led to anhedonia, we performed a sucrose preference test. Furthermore, a recent study indicated that BLA^CRF^ **→** NAc alters appetitive behaviors in a sex dependent fashion (18). Thus, we initially segregated mice by sex. (D) Sucrose preference across three days of testing in control and CRF cKO males (N = 4 and 7, respectively). There was no difference between groups (main effect of day: F_2,18_ = 0.1566, p = 0.8562; group: F_1,9_ = 0.01758, p = 0.8974; day x group: F_2,18_ = 0.2506, p = 0.7810). (E) Sucrose preference in control and CRF cKO females (N = 4 and 6, respectively). There was no difference between groups (main effect of day: F_2,16_ = 0.0582, p = 0.944; group: F_1,8_ = 0.0885, p = 0.7736; day x group: F_2,16_ = 0.9802, p = 0.3967). (F) Sucrose preference in control (black, N = 8) and CRF cKO (green, N = 13) mice across 3 days of testing. There was no significant difference between groups (main effect of day: F_2,38_ = 0.03851, p = 0.9623; group: F_1,19_ = 0.1361, p = 0.7162; day x group: F_2,38_ = 0.1047, p = 0.9008; N = 8 control, 13 cKO). (G) Total daily liquid intake for control and CRF cKO mice over 3 days of sucrose preference testing. There was no difference between groups (main effect of day: F_2,38_ = 0.8714, p = 0.4266; group: F_1,19_ = 0.04711, p = 0.8305; day x group: F_2,38_ = 1.114, p = 0.3389). (H) Representative locomotion traces from video monitored control and CRF cKO mice during the first minute in a novel open field test. (I) Distance traveled by control and CRF cKO mice in the novel open field test, binned by 1 min increments. (J) Total distance traveled by control and CRF cKO mice during the “novelty exploration” phase of the novel open field test (defined as the first minute). Control mice traveled significantly more than CRF cKO mice during this first minute (unpaired t-test, p = 0.0120, N = 9 each). (K) Average distance traveled by control and CRF cKO mice during the novel open field “sustained locomotion” period (defined as the last 10 m of the test). During this period, there were no significant locomotor differences between control and CRF cKO mice (unpaired t-test, p = 0.6725, n = 9 each) (Data are represented as mean ± SEM. Scale bar = 10 µm) (^ns^p > 0.05, *p < 0.05, **p < 0.01, two-way repeated measures ANOVA with Sidak’s post-hoc unless otherwise noted above.)

**Figure S11, related to Figure 6.**
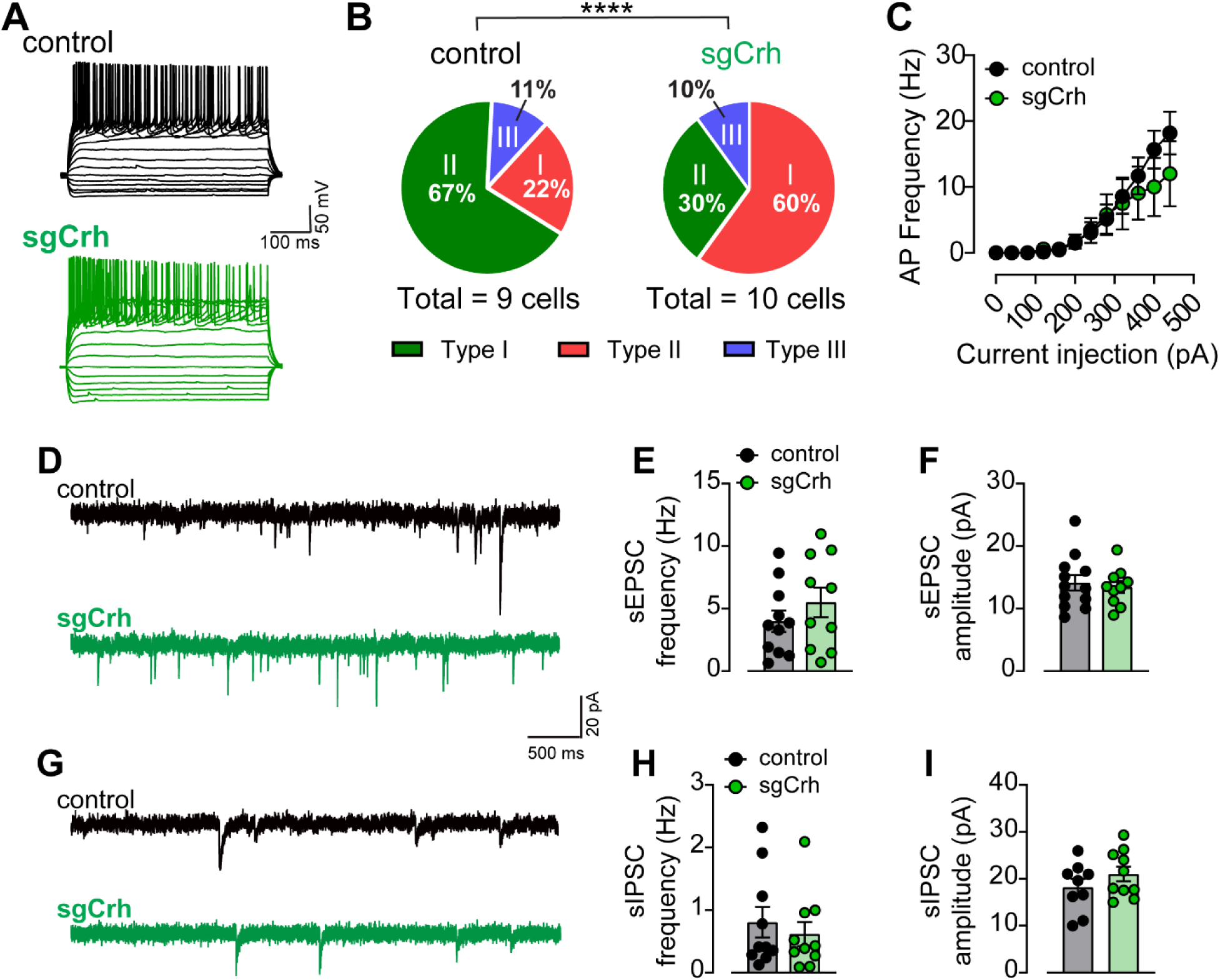
Deletion of CRF does not change electrophysiological properties of NAc^CRF^ cells. (A) Representative electrophysiological traces of firing from YFP+ cells from control sg*Rosa26* (black) and sgCrh (green) mice in response to current injections (-160 to +480, 40pA steps). (B) Distribution of CRF+ subtypes from control and sgCrh recordings. Cells from controls were primarily Type II, while cells from sgCrh mice were primarily Type I (Chi-squared test, p < 0.0001). (C) Input-output curve of firing in response to current injections from control and sgCrh current-clamp recordings. There was no significant difference in firing patterns between groups (Two-way RM ANOVA, current step x viral vector, F_11,176_ = 0.7829, p = 0.6568, N_Cells_ = 9 each) (D) Representative traces of spontaneous EPSCs (sEPSCs) from sgCrh and control mice. (E) Relative frequency and (F) amplitude of sEPSCs from control and sgCrh recordings. There was no difference between groups in either measure (unpaired t-tests, ps > 0.05). (G) Example traces from spontaneous IPSC (sIPSC) recordings from control and sgCrh cells. (H) Quantification of sIPSC frequency and (I) amplitude revealed no difference between control and sgCrh cells (unpaired t-tests, ps > 0.05). (Data are represented as mean ± SEM.)

**Figure S12, related to Figure 7.**
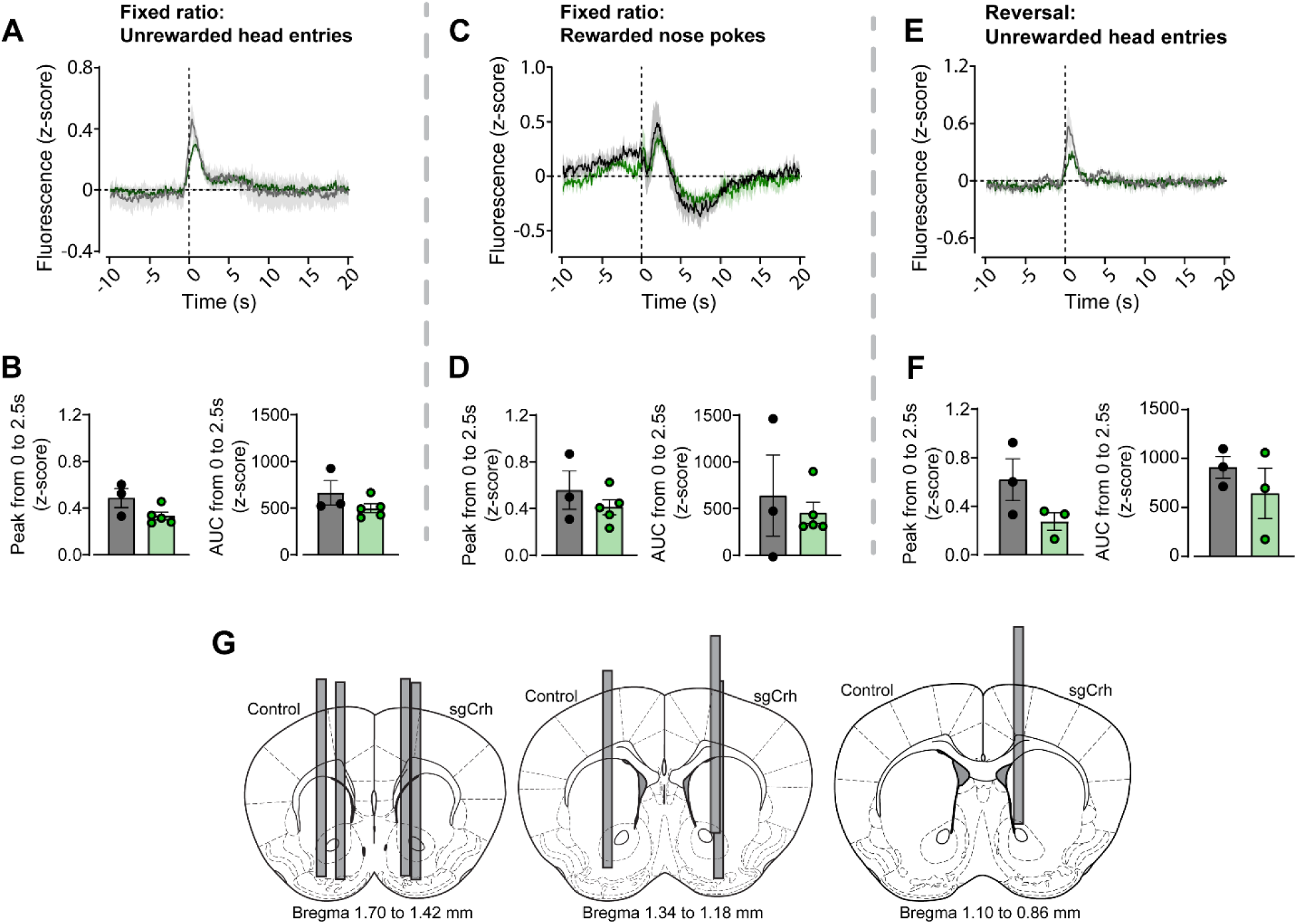
Additional fiber photometry data from CRF CRISPR experiments. (A-F) Average fluorescence (z-score) was aligned to behavioral event TTLs (at t=0) from control (black) and sg*Crh* (green) mice. The peak and AUC were calculated for 0 to 2.5 s following TTL. (A) Fluorescence following unrewarded head entries in fixed ratio task. (B) There was no difference between groups in peak (left) or AUC (right) in response to fixed ratio unrewarded head entries (C) Fluorescence following a rewarded poke in fixed ratio task. (D) Peak (left) and AUC (right) of fluorescence from fixed ratio task, time-locked to rewarded nose poke. There was no difference between groups. (E) Average fluorescence following an unrewarded head entry during the reversal test days. (F) Peak (left) and AUC (right) of fluorescence following an unrewarded head entry during reversal sessions. There was no significant difference between groups. (G) Schematic of fiber placements from all control (left hemisphere) and sgCrh (right hemisphere) mice included in CRISPR fiber photometry analysis. (Data are represented as mean ± SEM.) (p > 0.05, unpaired t-tests.)

